# Evolutionary turnover of key amino acids explains conservation of function without conservation of sequence in transcriptional activation domains

**DOI:** 10.1101/2024.12.03.626510

**Authors:** Claire LeBlanc, Jordan Stefani, Melvin Soriano, Angelica Lam, Marissa A. Zintel, Sanjana R. Kotha, Emily Chase, Giovani Pimentel-Solorio, Aditya Vunnum, Katherine Flug, Aaron Fultineer, Niklas Hummel, Max V. Staller

## Abstract

Protein function is canonically believed to be more conserved than amino acid sequence, but this idea is only well supported in folded domains, where highly diverged sequences can fold into equivalent 3D structures with identical function. Intrinsically disordered protein regions (IDRs) often experience rapid amino acid sequence divergence, but because they do not fold into stable 3D structures, it remains unknown when and how function is conserved. As a model system for studying the evolution of IDRs, we examined transcriptional activation domains, the regions of transcription factors that bind to coactivator complexes. We systematically identified activation domains on 502 homologs of the transcriptional activator Gcn4 spanning 600 MY of fungal evolution in the Ascomycota. We find that the central activation domain shows strong conservation of function without conservation of sequence. We identify the molecular mechanism for this conservation of function without conservation of sequence: evolutionary turnover (gain and loss) of acidic and aromatic residues that are important for function. We further see turnover of complete N-terminal activation domains. This turnover at two length scales confounds multiple sequence alignments, explaining why traditional comparative genomics cannot detect functional conservation of activation domains. Evolutionary turnover of key residues is likely a general mechanism for conservation of function without conservation of sequence in IDRs.

## Introduction

The evolution of eukaryotic transcription factors (TF) contains a paradox: TF protein sequences diverge quickly but maintain function over long evolutionary distances. For example, the master regulator of eye development in mice, Pax6, induces ectopic eyes in flies, and fly Pax6 (*eyeless*) creates ectopic eye structures in frogs and mice^1–3^. While the DNA-binding domains (DBD) are 96% identical, eye induction requires the intrinsically disordered regions (IDRs), which are only 35.5% identical. Despite substantial sequence divergence, these IDRs must share a conserved function. For folded domains, highly diverged sequences can fold into the same 3D structure, providing a molecular mechanism for conservation of function without conservation of sequence^4–6^. Here, we seek an analogous framework for understanding the evolution of IDRs. Small-scale studies have found examples of diverged IDRs that conserve function^7–9^ or do not conserve function^10,11^. Transcriptional activation domains provide an excellent model system for studying IDR evolution because they are one of the oldest classes of functional IDRs^12^, they are required for TF function, and their activity can be measured in high throughput^13^. Our goal is to identify molecular mechanisms by which TF IDR function can be conserved in the face of rapid sequence divergence.

Evolutionary studies of acidic activation domains in yeast benefit from high-throughput data that define sequence features controlling their function^13–19^. These data have trained neural network models for predicting activation domains from protein sequence^14,17,19–22^. Our acidic exposure model further provides a biophysical mechanism for the observed features: aromatic and leucine residues make key contacts with hydrophobic surfaces of coactivator complexes, but these residues can also interact with each other and drive collapse into an inactive state^13,23–26^. The acidic residues repel each other, expand the activation domain, and promote exposure of the hydrophobic residues. In many cases, the aromatic and leucine residues are arranged into short linear motifs. Large-scale mutagenesis showed the acidic exposure model applies to hundreds of human activation domains^27^.

We test the hypothesis that conservation of function without conservation of sequence in TF IDRs results from evolutionary turnover of functional elements. Evolutionary turnover is repeated gain and loss of functional elements. Mutations create new functional elements, and negative selection maintains a minimum number of elements, allowing ancestral elements to be lost. The combination of turnover and neutral drift gives the appearance that functional elements move around. For TFs, it is unclear if the functional elements will be entire activation domains, short linear interaction motifs (SLiMs)^28^, or individual amino acids. Here, we aim to identify the functional units and test whether evolutionary turnover can maintain function despite sequence divergence.

We investigated the molecular mechanisms by which full-length TFs can maintain activator function over long evolutionary distances despite divergence of their amino acid sequences. As a model system, we used 502 diverse homologs of Gcn4, a nutrient stress TF, and screened for activation domains with a high-throughput functional assay in *Saccharomyces cerevisiae*^*13*^. All homologs contain at least one activation domain. In the central activation domain, there is conservation of function without conservation of sequence across 600 million years of evolution. This functional conservation emerges from the evolutionary turnover of individual acidic and hydrophobic residues. In addition, we see evolutionary turnover of entire N-terminal activation domains. Our functional screening reveals that conservation of function without conservation of sequence in activation domains arises from turnover of individual residues.

## Results

### Characterization of a tiling-library of Gcn4 homologs

To study the evolutionary dynamics of TFs, we experimentally mapped activation domains across a diverse collection of homologous TFs. We and others have shown that protein fusion libraries, designed to tile across protein sequences with short, 30-60 amino acid peptides, can faithfully measure activation domain activity^13–15,17,18,29^. Furthermore, because activation domain function in yeast is a reliable measure of endogenous function in humans^30^, viruses^31^, *Drosophila*^*32,33*^, plants^19,34,35^, and other yeast species^36^, we reasoned that the activity of fungal homologs in our assay would serve as a faithful measure of activity in their native context. In all subsequent analysis, we assume that tile activity measured in *S. cerevisiae* is a good proxy for TF function in their native species.

As a null hypothesis, we assumed the TF function is conserved and that the observed diversity of sequence is the result of mutations, purifying (negative) selection, and neutral drift. We found evidence for weak negative selection on the full-length TF using a high-quality set of thirty-six Gcn4 homologs from the yeast gene order browser (**Figure 1A**)^37^. It follows that most of the sequence differences in this family are neutral.

**Figure 1:**
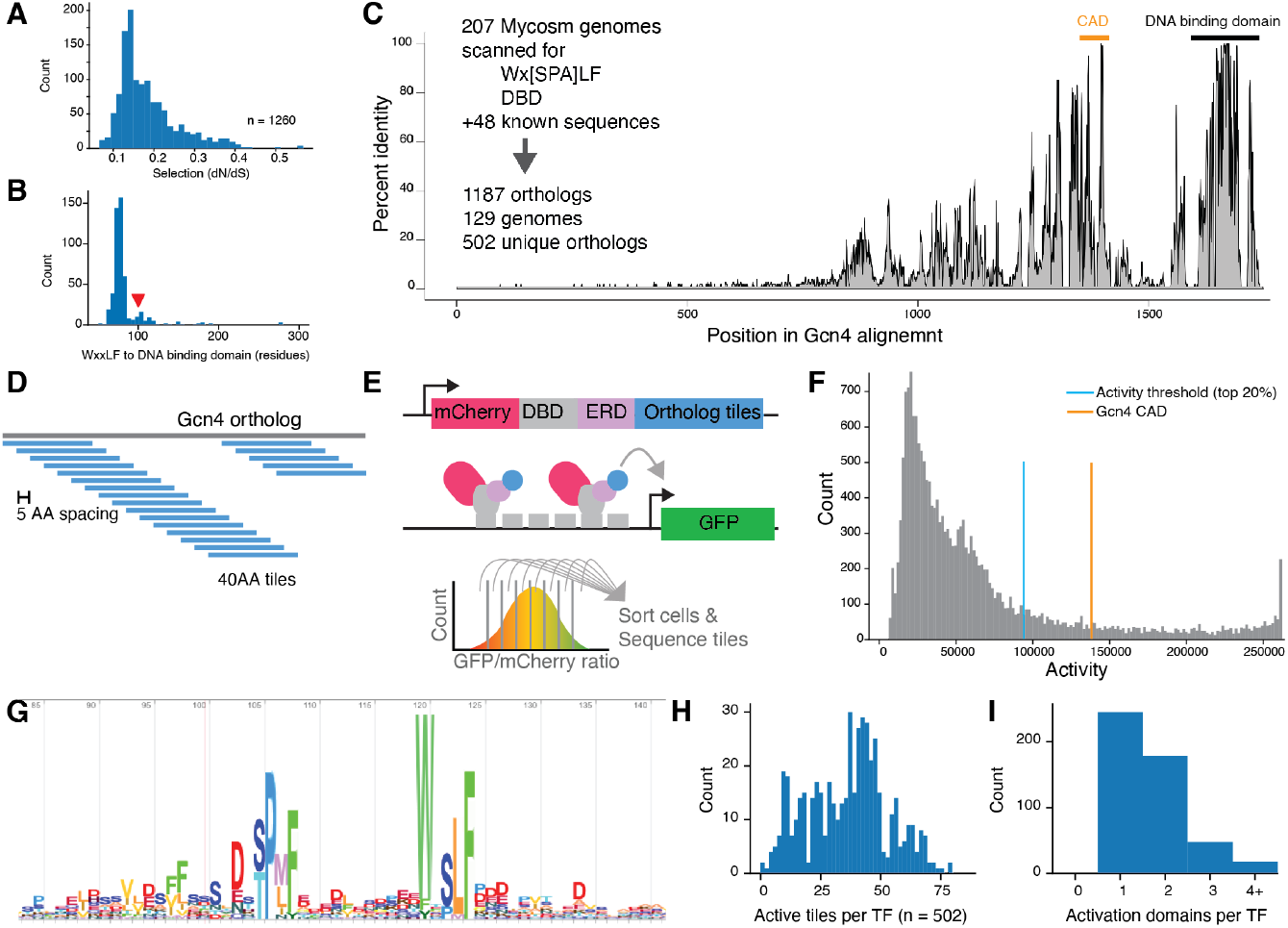
Screening fragments of Gcn4 homologs for activation domain activity in *S. cerevisiae*. A) The pairwise omega coefficients from PAML^39^ (dN/dS) from YGOB are all less than one, indicating negative selection. B) The distance between the WxxLF motif and the start of the DBD is conserved. Red arrow, *S. cerevisiae*. C) The MSA of 500 homologs shows the DBD binding domain is highly conserved, and the Central Activation Domain around the WxxLF motif is moderately conserved. Sequence divergence is driven by insertions: 54% of columns in the MSA contain fewer than 1% of sequences and are not shown in this MSA. D) The homolog tiling strategy and the high-throughput activation domain assay. E) The high-throughput assay for measuring activation domain function uses a synthetic TF with mCherry for quantification of abundance, the Zif268 DNA binding domain (DBD), an estrogen response domain (ERD) for inducible activation, and a C-terminally fused tile. Tile activity was calculated based on barcode abundance in eight equally sized bins of a FACS sorting experiment. Bins were set based on GFP/mCherry ratios. F) The distribution of measured tile activities with the activity threshold (top 20%). The *S. cerevisiae* Gcn4 central activation domain activity is shown in orange. G) The sequence logo from the 4th iteration of a search for Gcn4 homologs in fungal genomes with HMMER. This independent analysis confirmed the WxxLF motif is more conserved than the FF and MFxYxxL motifs. H) The number of active tiles found on each full-length TF (tiles that map to multiple homologs can count multiple times in this analysis). I) Combining overlapping active tiles shows that most TFs have two or more activation domains.

We found 502 unique Gcn4 homolog sequences from 129 genomes that span the Ascomycota, the largest phylum of Fungi, representing >600 million years of evolution^38^ (**Figure S1**). While the Gcn4 homologs vary in length (**Figure S1E**), 500 have the DBD at the C-terminus, and the distance between the WxxLF motif and the DBD is very consistent (**Figure 1B**).

The Gcn4 multiple sequence alignment (MSA) typifies eukaryotic TF evolution, with a highly conserved DBD and lower conservation in the rest of the protein (**Figure 1C)**. Distant pairs of sequences do not align outside of the DBD. The central activation domain (central activation domain) shows intermediate levels of conservation, driven in part by the WxxLF motif (**Figure 1C, G**).

### High-throughput measurement of activation domains on the homolog

To study the evolution of TF function, we mapped activation domains on the homologs. For each of the 502 Gcn4 homologs, we tiled across the full-length protein with 40 amino acid (AA) tiles spaced every 5 AA and measured activities of all tiles in *S. cerevisiae* using a high-throughput assay^13^ (**Figure 1D, 1E**). We recovered 18947 of 20731 designed tiles (91.4%), and these data were of high quality (Methods, **Figure S2, S3**). As a threshold for highly-active tiles, we used the top 20% of sequences. When we combined overlapping active tiles, most TFs had multiple activation domains (**Figure 1H, 1I)**. Due to the divergence of the homologs, the sequences of the active tiles are very diverse, allowing us to study sequence-to-function relationships controlling activation domain function. To our knowledge, this dataset is the largest functional study of TF evolution to date.

### Conservation of function without conservation of sequence in the Central Activation Domain

The central activation domain of Gcn4 is functionally conserved across the homologs. We used overlapping tiles to infer the activity of each position in each full-length protein (**Figure 2**, Methods). We found that nearly all homologs had high activity in the central region. This central activation domain can drift side-to-side but stays near the WxxLF motif (**Figure S4, S5, S6**). The sequences of this central region are highly diverged, indicating conservation of function without conservation of sequence (**Figure S7**).

**Figure 2:**
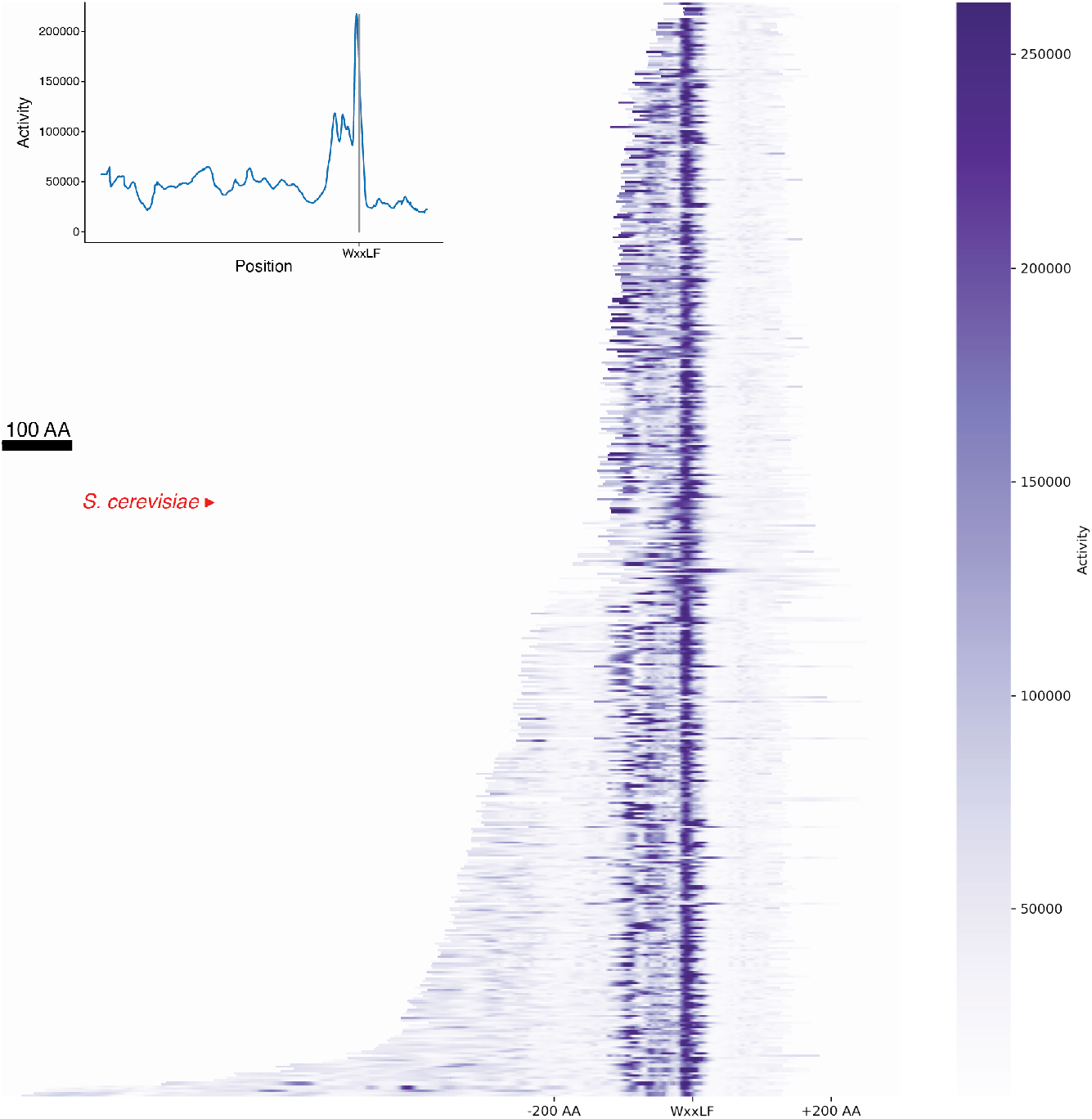
The central acidic activation domain of Gcn4 is functionally conserved. We used the tile activity data to impute the activity of each position in all the homologs and visualized these activities as a heatmap. Four-hundred ninety-seven homologs are sorted by length and aligned on the WxxLF motif. Inset, vertically averaging the heatmap indicated the peak is ten residues upstream of the WxxLF motif. Activity is consistently high around the WxxLF motif, indicating deep functional conservation. Upstream activity is more salt and pepper, indicating recurrent gain and loss of N-terminal activation domains. Red arrow, *S. cerevisiae*. Black scale bar, 100 AA.

We sought to understand the molecular mechanism behind this conservation of function without conservation of sequence. Identifying the amino acids that control the function of individual tiles allowed us to rule out turnover of short linear motifs and demonstrate evolutionary turnover of key amino acids.

### The sequence features of highly active tiles indicate a flexible grammar

The Gcn4 homolog dataset contains all previously observed relationships between sequence and function (**Figure S8**). Active tiles contain many acidic residues and WFYL residues (**Figure 3A, S9**) consistent with the acidic exposure model (**Figure 3B, 3C**). In the control activation domains, all published motifs of aromatic and leucine residues made large contributions to activity, but no individual motif was sufficient for full activity (**Figure S3**).

**Figure 3:**
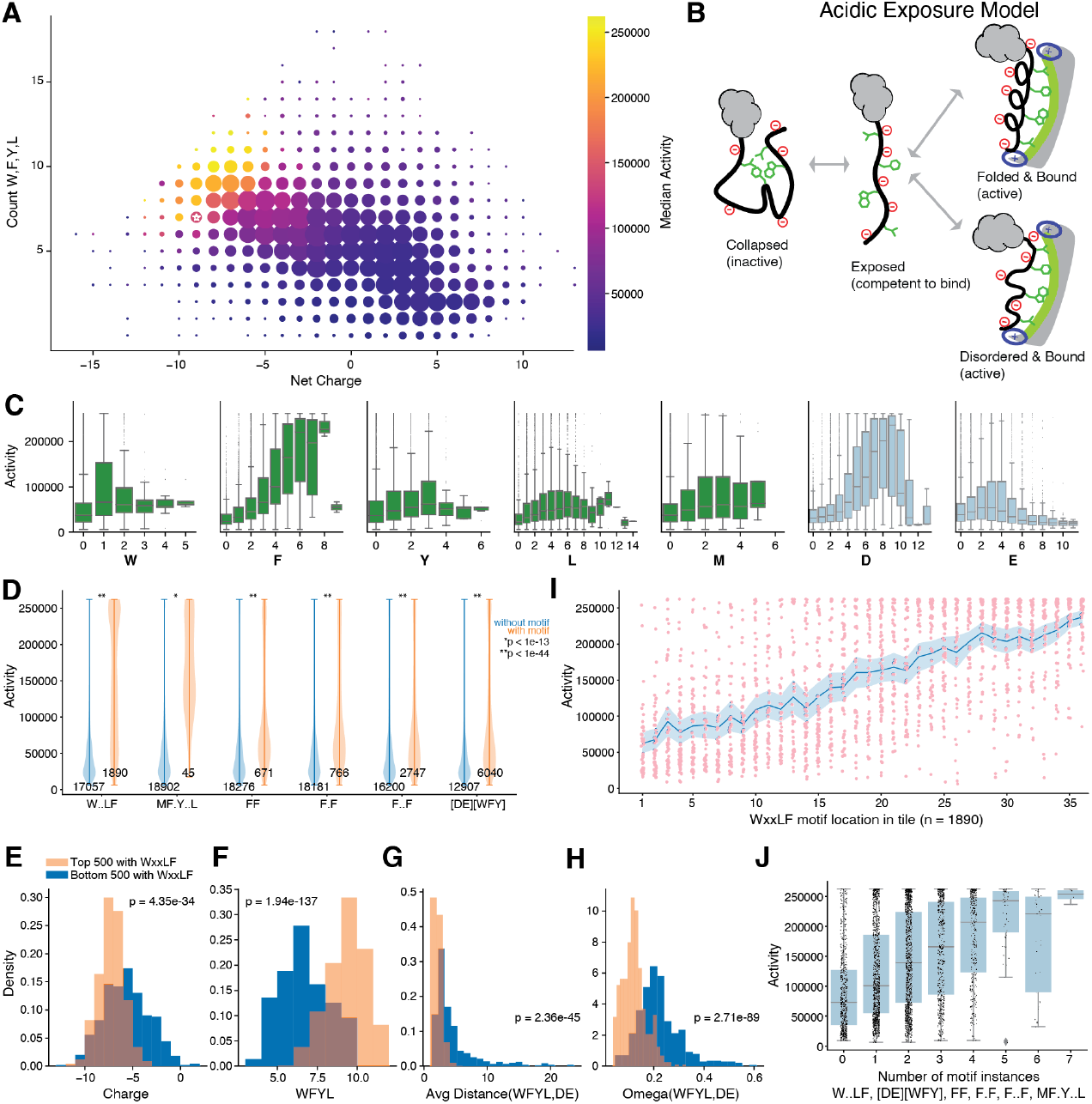
Highly active tiles contain acidic, aromatic, and leucine residues, supporting the acid exposure model of acidic activation domain function. **A**) For each tile, we compute net charge and count the number of WFYL residues. The area of the point indicates the number of tiles with the combination of properties. The color is the median activity. White star, *S. cerevisiae* Gcn4. **B**) The acidic exposure model of acidic activation domain function. **C**) Box and whisker plots for the residues that make the largest contributions to activity. Gray points are outliers beyond the whiskers (1.5x the interquartile range). **D**) For each published motif, tiles that contain the motif (orange) have higher activity than tiles without the motif (blue). p-values are uncorrected Welch’s t-test. E-H) Comparison of the 500 most active tiles with the WxxLF motif to the 500 least active tiles with the WxxLF motif. The active tiles had more acidic residues (E), more W,F,Y,L residues (F), closer spacing between WFYL and acidic residues (G), and more intermixing as measured by Omega^40,41^ (H). **I**) For each tile with the WxxLF motif, activity is plotted against the location of the W. Blue, mean and 95% confidence interval. The location of the motif is correlated with activity. **J**) Tiles with more published motifs are more active.

We searched for sequence grammar, the arrangement of amino acids, that controls activation domain function. We and others have previously observed that acidic activation domains use a highly flexible grammar^13,14,17,24,42^. As a baseline, we quantified how amino acid composition contributes to function with ordinary least squares (OLS) regression on single amino acids, which explains 49.9% of variance in activity (**Table 1**, Area under the receiver operator characteristic (AUC) = 0.9346, and area under the precision recall curve (PRC) = 0.7620, **Table S9**). Regression on dipeptides^17^ led to sixty-nine significant parameters that explain 60.2% of the variance in activity (**Table 1**, AUC = 0.9472, PRC = 0.8190). More complex sequence motifs did not improve the regression models: published motifs explained 33.1%, and forty *de novo* motifs explained 50.5% of the variance in activity (**Table 1**). Composition and dipeptides are major determinants of activity, consistent with weak grammar.

**Table 1:** Ordinary Least Squares regression on tile composition explains a large fraction of the variance in measured activation domain activity

| Model | Number parameters | Number of statistically significant parameters | Adjusted R<sup>2</sup> |
| --- | --- | --- | --- |
| Single AAs | 20 | 16 | .498 |
| Single AAs - reduced | 16 |  | .498 |
| Dipeptides | 400 | 69 | .651 |
| Dipeptides - reduced | 69 |  | .608 |
| Published Motifs | 7 | 5 | .334 |
| *de novo* motifs | 40 | 27 | .502 |
| *de novo* motifs - reduced | 27 |  | .500 |
| *de novo* motifs + single AAs | 60 | 37 | .606 |
| *de novo* motifs + single AAs | 37 |  | .604 |

To further search for grammar, we compared tiles with the WxxLF motif that had high or low activity (**Figure 3D**). Highly active tiles were more acidic (**Figure 3E**) and had more WFYL residues (**Figure 3F**). The first grammar signal is that active tiles have shorter distances between DE and WFL residues (**Figure 3G**). The second grammar signal is that tiles with more evenly intermixed acidic and W,F,Y,L residues are more active, supporting the acidic exposure model^41^ (**Figure 3H**, Methods). The strongest grammar signal is that tiles with the WxxLF motif near the C-terminus are more active (**Figure 3I**). The additional negative charge of the C-terminus likely increases exposure of the motif. This result emphasizes how a conserved short linear motif requires both an acidic context and supporting hydrophobic residues to create an activation domain. Tiles with more published motifs tend to be more active, consistent with multivalent binding to coactivators^17,43^ (**Figure 3J**).

In conclusion, these yeast activation domains are nucleated by aromatic residues (W,F,Y) and supported by L and M residues. These hydrophobic residues require an acidic context (D and E residues).

### Turnover of key amino acids facilitates conservation of function without conservation of sequence in the Central Activation Domain of Gcn4

The central activation domain region of the Gcn4 homologs showed strong conservation of function without conservation of sequence. We investigated whether this phenomenon could be explained by evolutionary turnover of motifs or evolutionary turnover of individual residues.

We found no evidence for turnover of motifs that control activation domain function. Each of the published motifs contributed to activity (WxxLF, MFxYxxL, FF, FxF, FxxF, and [DE][WFY] **Figure S3**) and was enriched in active tiles (**Figure 3D**), but only the WxxLF motif was conserved (**Figure 1G**). We did not detect the emergence of new instances of these motifs, so we can reject the motif turnover hypothesis. We see conservation SP motifs, but these motifs are not involved in activation domain strength. The full-length homologs contain up to four SP motifs upstream of the WxxLF motif. In *S. cerevisiae*, this SP is a TP (T105), which is phosphorylated to create a phosphodegron that shuts down the Gcn4 program during the recovery from starvation^44,45^. Conservation and turnover of SP motifs may contribute to regulation of protein degradation.

We found evolutionary turnover of individual phenylalanine (F) residues that are critical for high activity (**Figure 4, S10**). We selected a seventy residue region around the WxxLF motif (W-50 : W+19) that contained the peak of inferred activity. We examined the sequences of the 50% most active sequences. There are only two positions where an F is present in the majority of sequences but a total of ten positions where F is the most common residue, a signature of turnover. The key F residues experience evolutionary turnover, giving the appearance of moving around the activation domain.

**Figure 4:**
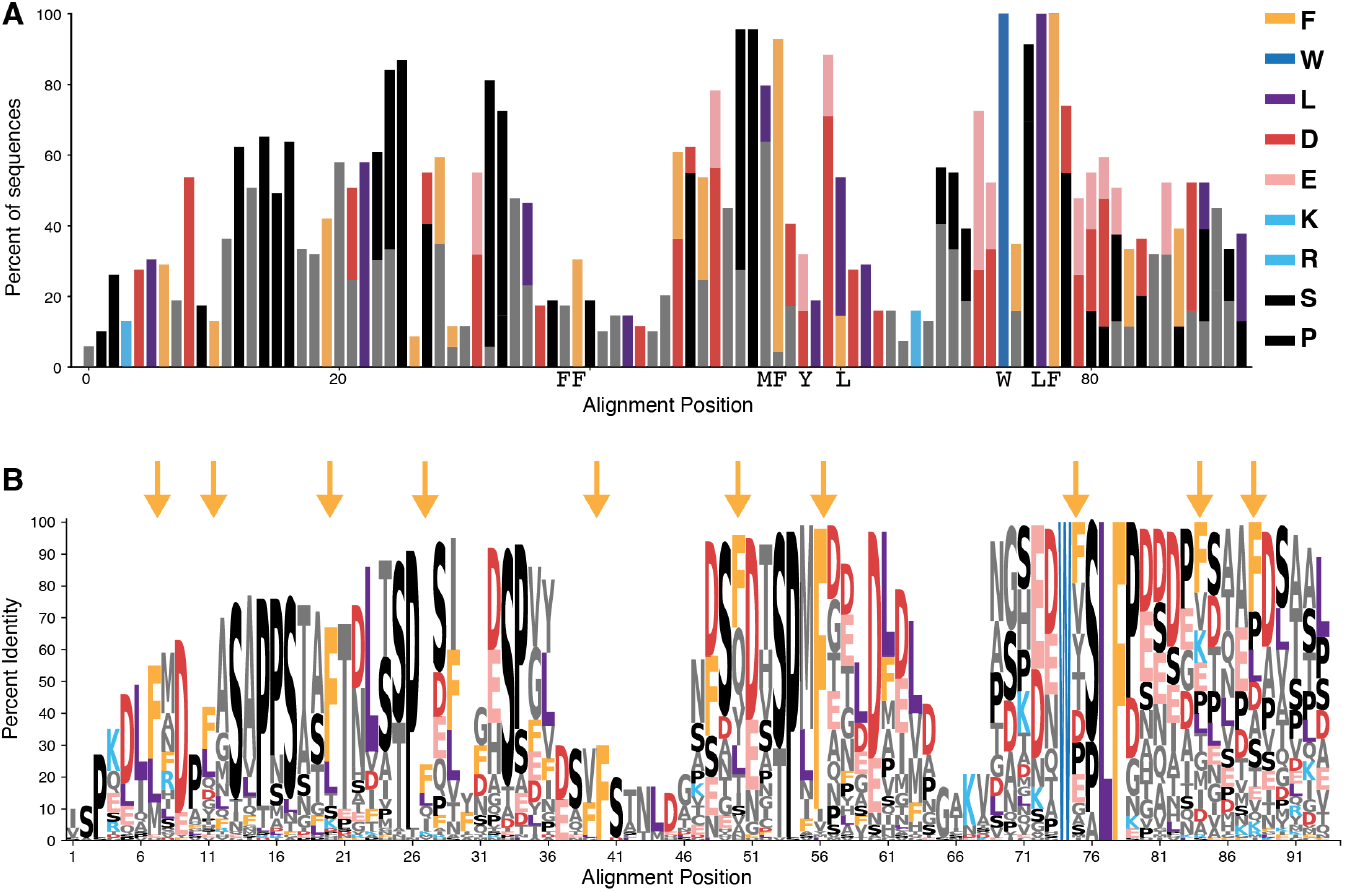
Evolutionary turnover of aromatic and acidic residues explains the conservation of function without conservation of sequence in the central activation domain of Gcn4. **A**) For the sixty-nine most active unique regions around the WxxLF motif, a bar plot showing the relative amino acid frequencies from the MSA. The acidic residues, D and E, interchange. **B**) A sequence logo for the MSA. Arrows indicate the ten positions where F is the most abundant residue. There is some interchange between F and L. Black, SP motifs.

In addition, we observe evolutionary turnover of acidic residues within the central activation domain. In the highly active sequences, individual acidic positions (D and E) interconvert (**Figure 4A,B, S10**). When pooled together, D+E conservation matches or exceeds the conservation level of many aromatic residues. Acidic residues interconvert and move around the activation domain, hallmarks of turnover.

### Additional support for turnover of individual amino acids

To this point, all of our analysis has used only the MSA, so we next leveraged the additional information present in the species tree. We tested the hypothesis that the gains of F residues precede the loss of F residues. In most cases, there is too much evolutionary distance between the species to answer this question. However, in the high-quality YGOB alignment^37^, we see the gain of an F precedes the loss of an ancestral F (**Figure 5A**). This example of gains preceding loss bolster the evidence for evolutionary turnover of key residues.

**Figure 5:**
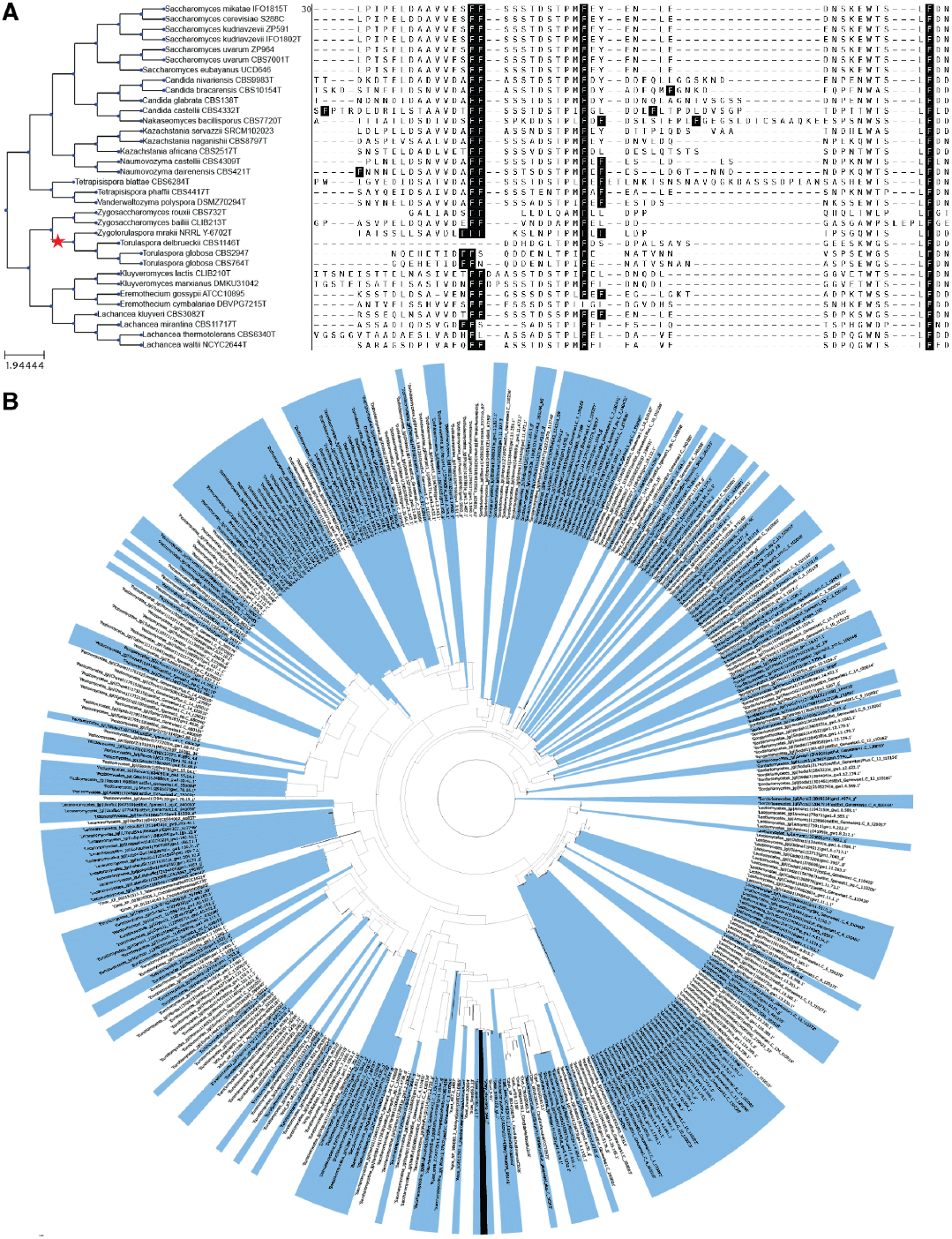
Turnover of phenylalanine residues and recurrent loss of N-terminal activation domains. A) In the YGOB MSA, 34/36 species have xFF. However, *Zygotorulaspora mrakii* has FFF, the adjacent *Torulaspora globosa CBS2947* has FFN and *Torulaspora globosa CBS764T* has FFS. This pattern is consistent with the local ancestor gaining a third F, one descendent keeping all three, and two descendents losing one ancestral residue to revert to 2 Fs. All the extant sequences have high activity. Red star, proposed gain of 3rd F. Tree visualization was made with ETEToolkit^5^. B) Homologs that contain N-terminal, non-WxxLF activation domains are highlighted in blue on the gene tree. This pattern is consistent with recurrent gain and loss of N-terminal activation domains.

The turnover and conservation patterns we observed in the Gcn4 homologs generalized to other activation domains. For four TFs, we searched for homologs in the Y1000+ collection and made alignments of their activation domains (**Figure S11**). In these MSAs, aromatic residues were highly conserved and acidic residues interchange at many positions. Some positions also showed interchange between aromatic residues. Other regions showed turnover of aromatic and leucine residues. When we reanalyze measurements of Pdr1 homologs, we see similar patterns^14^. We conclude that evolutionary turnover of aromatic, leucine, and acidic residues enables conservation of function without conservation of sequence in acidic activation domains.

### Turnover of entire activation domains

At the protein level, we found turnover of entire activation domains. *A priori*, it was not a given that all the Gcn4 homologs would be activators, because on long evolutionary timescales, a family of TFs that share a conserved DBD will include both activators and repressors ^19,27,29,35^. All homologs had at least one tile that functioned as an activation domain in our assay **(Figure 1H**), indicating that activator function is conserved across 600 million years of evolution. The one exception is a deprecated gene model, and the longer model of this locus has an activation domain (Methods). After combining overlapping active tiles, the majority of homologs have more than one activation domain (**Figure 1I**). Virtually all the additional activation domains are N-terminal, and their distribution across the phylogeny is consistent with recurrent gain and loss (**Figure 5**). These N-terminal activation domains show intermediate conservation in the MSA (**Figure S5**), and their sequences are diverse (**Figure S6**). Recent insertions are depleted for activity (**Figure S6B**). Together, these data demonstrate turnover of entire activation domains.

### Machine learning models support evolutionary turnover

The Gcn4 homologs provide a large, unique dataset to evaluate deep learning models that predict activation domains from amino acid sequence. We compared two first-generation neural networks^14,17^ with two second-generation models that we developed^19,20^. All the models can approximate the locations of activation domains in full-length TFs, but the new model, TADA, is substantially more accurate at predicting the activities of individual tiles and identifying activation domain boundaries (**Figure S12**).

We used TADA to predict the contributions of F residues to activity in the central activation domain. It predicted that all the F residues contribute to activity (**Figure 6A, 6B, 6C**). Importantly, contributions of the most conserved F positions are indistinguishable from recently evolved F positions (**Figure 6D**). This analysis further supports evolutionary turnover of key F residues.

**Figure 6:**
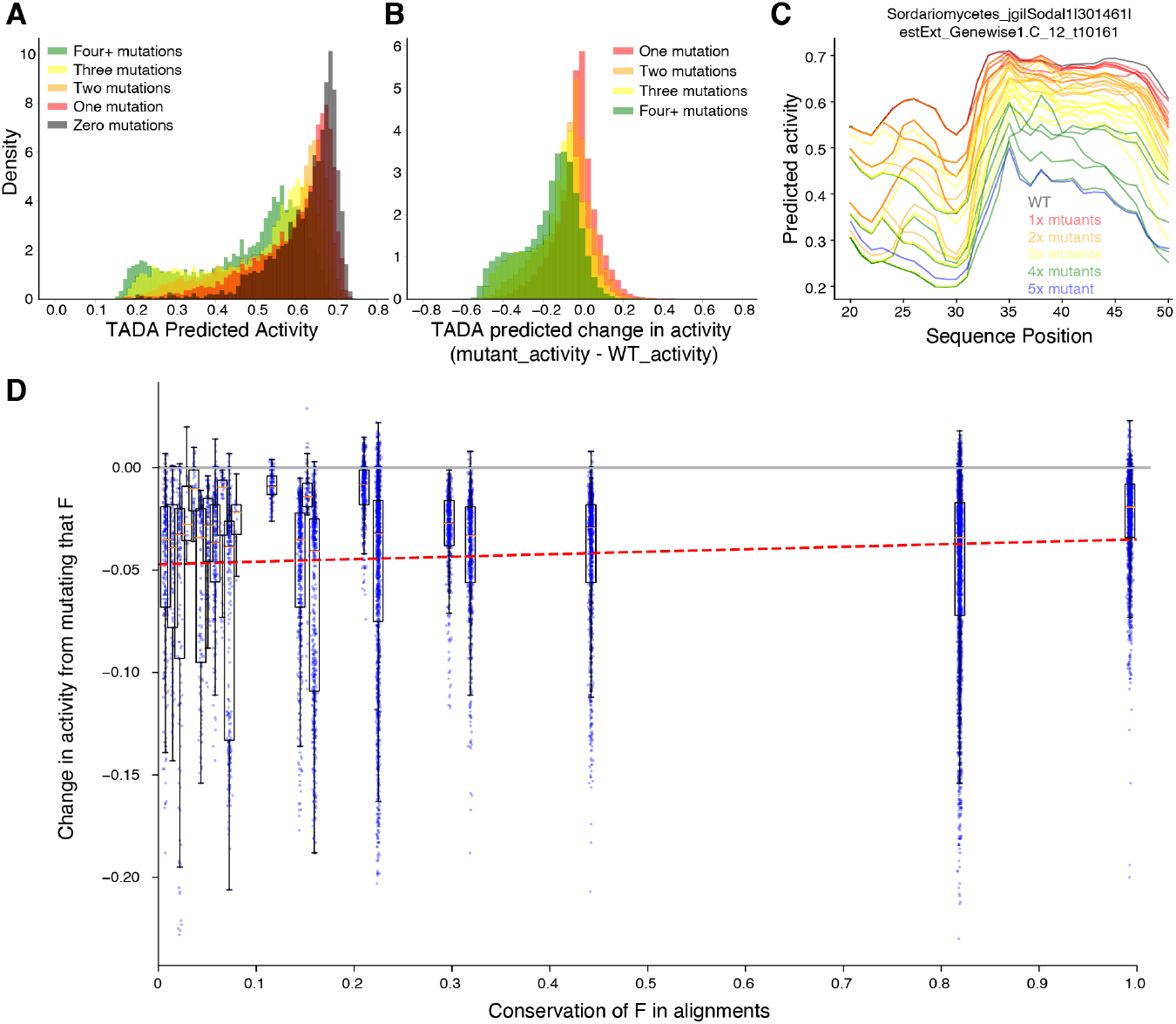
Predicted activity of phenylalanine mutations supports evolutionary turnover. A) Starting with the 138 unique 70 AA regions around the WxxLF motif, we tiled these regions into 40 AA tiles spaced at 1 AA. For each resulting tile, we predicted activity (gray) with TADA. Next we generated all single phenylalanine to alanine (F>A) mutations and predicted activity (red). We repeated this process for double (orange), triple (yellow), and 4+ (green) F>A mutations. Tiles with more mutations generally had lower activity. B) We calculated the change in predicted activity for the mutations in A. Nearly all mutations decreased predicted activity, consistent with earlier analysis. C) For a single representative region, the traces indicate predicted activity of the WT (black), all single F mutants (red), double F mutants (orange), triple F mutants (yellow), four F mutants (green) and five F mutants (blue). Mutations decrease predicted activity. D) Using only the single F>A mutations, we grouped mutants by the observed conservation of the F residue in the 138 sequences. Most mutations decrease activity (they are below the pink line). More conserved F residues generally do not cause larger predicted decreases in activity when mutated, because the regression line (red) is flat. This analysis suggests all the F’s contribute to activity. This result supports the turnover of key F residues.

## Discussion

By functionally screening protein fragments from a family of homologous TFs, we identify two molecular mechanisms for how TFs show strong conservation of function without conservation of sequence: evolutionary turnover of complete activation domains and evolutionary turnover of key acidic and phenylalanine residues within activation domains. Our results emphasize how IDR function can be highly conserved and constrained yet invisible in traditional comparative genomics.

Evolutionary turnover of key residues arises from the weak grammar constraints on activation domains. Weak grammar explains why multiple screens for activation domains have found only one recurrent motif, LxxLL, which can be important for binding the Kix domain^14,17,27,28,46,47^. Here, we argue that yeast activation domains are nucleated by clusters of W and F residues surrounded by acidic residues and boosted by Y, L, and M residues. Under this weak molecular grammar, individual residues are easily replaced, facilitating turnover. When only a few sequences are examined, clusters look like motifs. Each TF family has a different conserved cluster of hydrophobic residues that represents a good solution to binding the preferred coactivator. As a result, each TF family will appear to have a conserved, essential motif, but convergent evolution of motifs is rare.

Evolutionary turnover of key residues is possible because of the physical flexibility of the Gcn4 Med15 interaction. This study adds an evolutionary dimension to the idea that the physical flexibility of IDR interactions with folded domains allows for multiple binding modes^46,48,49^. This physical flexibility of these interactions in turn allows evolutionary plasticity. The Gcn4 central activation domain undergoes coupled folding and binding with the Med15 activation domain binding domains, but this interaction is a physically flexible, fuzzy interaction^43,50,51^. The short helix presents the WxxLF motif in many orientations to Med15, and molecular dynamics simulations suggest that these orientations interconvert^52^. This binding interaction imposes few structural constraints on the Gcn4 central activation domain, enabling evolutionary plasticity. Binding one sequence in multiple orientations is a step towards binding diverse homologs, which in turn is a step towards binding to many activation domains^14,43,50,51,53,54^. Coactivators that impose weak structural constraints on activation domains can become engines for evolutionary diversification of activation domains through neutral drift, creating an enormous sequence reservoir for later selection. This diverse sequence reservoir allows for selection on standing variation in new environments, creating evolvability.

Our deep dive into the evolution of one family complements other studies of IDR evolution. Using small numbers of sequences, conservation of IDR function across homologs has been observed, but often the essential residues are unknown^7,55^. In other systems, there is functional conservation of diverged IDRs, but the key residues are conserved^9,56^ or motifs are conserved^57^. In other cases, functional conservation results from the composition, but not the arrangement, of residues through emergent properties like net charge^8,58–62^. These cases likely permit even more turnover than we observe in Gcn4. In Afb1, IDR function is not conserved^10^, and the Msn2/4 IDRs have two overlapping functions, only one of which is conserved^11^. The closest parallel to our turnover of key residues is *de novo* evolution of phosphorylation motifs^63^. There remains a need for better IDR-alignment algorithms or alignment-free methods to group functionally related IDRs.

Our results fit well with findings that at long evolutionary distances, transcriptional regulatory networks rewire, substituting individual TFs but maintaining circuit logic^36,64,65^. Here, we examined much longer evolutionary distances and found that all the Gcn4 homologs are activators, indicating that the signs of TF connections are more conserved than individual connections. Slow changes in TF function reduce pleiotropy and may make it easier to substitute TFs at individual regulatory elements.

The turnover of key hydrophobic residues in activation domain evolution bears strong parallels to the turnover of TF binding sites in enhancer evolution. Metazoan enhancers are regulatory DNA that contain clusters of TF binding sites^66^. The DNA sequence of enhancers diverges rapidly as individual TF binding sites are gained and lost while maintaining function^67–69^. Orthologous enhancers are often impossible to detect in sequence alignments but can be identified by searching for clusters of TF binding sites^70–72^. Turnover of entire activation domains on a full-length TF parallels turnover of entire enhancers in a locus^73^.

Within activation domains, turnover of key residues parallels turnover of TF binding sites within enhancers. Both activation domains^22,23^ and enhancers^74^ have very flexible grammar. Given that TFs function by binding to enhancers, it is striking that both the protein and the DNA are evolving in the same way. Turnover of enhancers and TF binding sites endows gene expression with robustness to environmental stress and evolutionary plasticity^75–79^. Turnover of key residues in activation domains may similarly endow TFs with plasticity and robustness.

The primary limitation of this work is that we measured the activities of short fragments in one species. Measuring short, uniform fragments makes the experiments possible but can miss longer emergent activation domains^51,80^. If, in some species, an activation domain and cognate coactivator together experience many compensatory mutations, the assay may not detect activity. Our analysis of Med15 coactivator conservation shows that the four activation domain binding domains are conserved (**Figure S13**). Activity of our reporter is well correlated with Med15 binding affinity *in vitro*^*14*^. The most active tiles are computationally predicted to bind Med15^81^ (**Figure S14**). In the future, limited screening in additional species or screening tiles of multiple tile lengths would enrich this work.

## Materials and Methods

### Identification of homologous sequences

We computationally screened for Gcn4 homologs of *S. cerevisiae*. We started with a hand-collected set of forty-nine homologs, forty-eight of which contained the WxxLF motif ^13,51^. To find new homologs, we used two criteria: the bZIP DNA binding domain (IPR004827) and the regular expression Wx[SPA]LF for the WxxLF motif. These criteria distinguished Gcn4 homologs from other leucine zipper DNA binding domain TFs. We scanned 207 diverse and representative proteomes from the MycoCosm database (mycocosm.jgi.doe.gov). This computational screen yielded 1188 gene models from 129 genomes. These 1188 gene models combine to yield 502 unique proteins (**Table S1, Figure S1**).). Of these, >99% were reciprocal Blast best hits with *S. cerevisiae* Gcn4. This initial analysis was performed in 2020 by Sumanth Mutte of MyGen Informatics. Eighty-four of the genomes were from MycoCosm, while the original homolog collection contributed forty-five species. Genomes contained 1-32 gene models and 1-11 unique protein sequences (**Figure S1**). These sequences span nearly all the Ascomycota, the largest phylum of Fungi, representing >600 million years of evolution ^38^. The 502 unique homologs have variable lengths (**Figure S1E**), but the DBD is at the C-terminus in 500 homologs, and the distance between the WxxLF motif and the DBD is very consistent (**Figure 1B**).

All species were from the Ascomycota except for five entries with three unique sequences from Blastocladiomycota (**Figure S1**). The Blastocladiomycota homologs are the only proteins where the WxxLF motif does not align in the MSA. The sequence context of their WxxLF motif is H-rich instead of acidic:

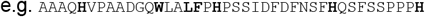

The Blastocladiomycota tiles with the WxxLF motif have high activity in the assay. The regions of Blastocladiomycota homologs that align to the WxxLF motif in the MSA have low activity in the assay. We suspect the N-terminal WxxLF in the Blastocladiomycota may have been gained by convergent evolution.

The Yeast Gene Order Browser has reconstructed the local synteny of the Gcn4 locus for thirty-seven genomes yielding a high-quality set of true homologs ^37^. Thirty-six species and the inferred ancestor contain one Gcn4 gene. *Kazachstania saulgeensis CLIB1764T* is missing a Gcn4 homolog. All of the post-whole-genome duplication species in this set contain only one Gcn4 homolog, suggesting there is no advantage of retaining two copies. All but one of the thirty-six homologs, *Zygosaccharomyces bailiii* ZYBA0L03268g, contain the WxxLF motif. Instead, *Z. bailiii* has an insertion in the WxxLF motif yielding W**PSL**EPLF. This sequence was not included in our experiment but was previously measured in a 44 AA tile, LDQAVVDEFFVNDDAPMFELDDGASGAWPSLEPLFGEDEERVAV and had high activity in Replicate 2 of our previous paper ^13^. This example further supports the observed conservation of function without conservation of sequence.

Despite substantial sequence divergence, all homologs show negative selection at the level of the full protein in the precomputed YGOB analysis. We downloaded a list of thirty-six pairwise Ka, Ks, and omega coefficients calculated from the yn00 output of Phylogenetic Analysis by Maximum Likelihood (PAML) (**Table S14**, November 2024).

We confirmed that the WxxLF motif is well conserved in fungal TFs with HMMER. We ran the web server for HMMER with default parameters, using *S*.*cerevisiae* Gcn4 as the seed sequence and restricting our search to Fungi. In the second, third, and fourth iterations of this search, the WxxLF motif was the most prominent feature of the profile HMM in the central region of the TF and always much more prominent than all other published motifs ^17,80^. **Figure 1G** shows the pHMM from the fourth iteration.

For the full-length homologs, MSAs were performed in Genious with the MAFFT algorithm (**Table S2**). We removed the two longest homologs that had the DBD near the center. In the MSA, 54% of positions had less than 1% identity and 88% had less than 5% identity.

Short alignments were created with MUSCLE online (https://www.ebi.ac.uk/Tools/msa/muscle/) or with or with MAFFT v7.526 and visualized with weblogo.berkeley.edu or the LogoMaker Python package.

### Design of the Gcn4 oligo library

We took the 502 unique protein sequences and computationally chopped them into 40 AA tiles spaced every 5 AA (e.g. 1-40, 6-45, 11-50 etc.). As a result, if two closely related sequences contain identical regions, insertions or alternatives (start sites) that change the phasing, a single tile can map to multiple full-length homologs. We removed duplicate tile sequences, yielding 20679 unique tiles. We added fifty-two control sequences (controls were included twice in the oligo pool to increase the probability they were recovered in the plasmid pool during cloning). The controls included hand-designed mutants in control activation domains and a handful of sequences from our previous study^13^ (**Table S3**, Control sequences). The final design file contained 20783 entries.

We reverse-translated tile sequences using *S. cerevisiae*-preferred codons. We added primer sequences for PCR amplification and HiFi cloning (‘ArrayDNA’ column in **Table S5**). We also added four Stop codons in three reading frames to ensure translational termination, even if there were one or two bp deletions, the most common synthesis errors. We used synonymous mutations to remove instances where the same base occurred four or more times in a row to reduce DNA synthesis errors. The resulting oligo pool was ordered from Agilent Technologies. The final oligos were of the form (see primer sequences in **Table S4**):

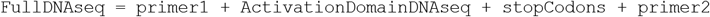

### Plasmid Library construction

The oligos were resuspended in 100 uL of water, yielding a 1 pM solution. The oligos were amplified with eight reactions of Q5 polymerase (NEB) using 1 ul of template, five cycles, Tm =72C, and the LC3.P1 and LC3.P2 primers. The eight reactions were combined into a single PCR clean-up column (NEB Monarch).

The backbone was prepared by digesting 16 ug of pMVS219 with NheI-HF, PacI, and AscI in eight reactions. We digested for seventeen hours at 37C and heat-inactivated for one hour at 80C. The desired 7025 bp fragment was run on a 0.8% gel, visualized with SYBR Safe (Invitrogen), and gel purified (NEB Monarch Kit). Note pMVS219 and pMVS142 have the same sequence, but the pMVS142 stock developed heteroplasmy, so we repurified it as pMVS219 and submitted the corrected stock to AddGene. Both pMVS219 and pMVS142 correspond to AddGene #99049.

We used NEB HiFi 2x mastermix to perform Gibson Isothermal Assembly to create the plasmid library. The 4x reaction volume had 328 ng of backbone and excess molar insert. We incubated at 50C for 15 min and assembled a backbone-only control in parallel. The assemblies were electroporated three times each into ElectroMax 10b E.coli (Invitrogen 18290-015) following the manufacturer’s protocol. A dilution series was plated and the bulk of the cells grown overnight in 140mL LB+Amp. These cultures overgrew, so they were spundown and frozen. The cultures were regrown with 105 mL LB+Amp and a MaxiPrep was performed (Zymo). An estimated 4.2 million colonies were collected, covering the library 200-fold.

To assess the quality of the plasmid library, we prepared an amplicon sequencing library (see below). Three independent amplicon libraries were prepared, and sequences present in all three were considered to be present in the plasmid pool with high confidence. GREP for the flanking NheI and AscI sites was used to pull out the designed fragments. Only perfect matches were used in this analysis. 20717 of 20731 designed sequences were detected (99.9%). The vast majority sequence abundances were within fourfold of each other, indicating minimal skew in library member abundance.

### Yeast transformation

The plasmid library was integrated into the DHY213 BY superhost strain, MATa his1Δ1 leu2Δ0 ura3Δ0 met15Δ0 MKT1(30G) RMEI(INS-308A) TAO3(1493Q), CAT5(91M), MIP(661T) SAL1+ HAP1+, a generous gift from Angela Chu and Joe Horecka. Requests for the parent strain are best directed to them. We integrated our library into the URA3 locus with a three-piece PCR ^82^. The upstream homology between URA3 and the ACT1 promoter was created by PCR amplifying the pMVS295 (Strader 6161) with the primers YP18 and CP19.P6. The downstream homology between the TEF terminator of KANMX and URA3 was amplified from pMVS196 (Strader 6768) with the primers YP7 and YP19. These template plasmids were a generous gift from Nick Morffy and Lucia Strader. To avoid PCR, the plasmid library was digested with Sal I-HF and EcoRI-HF (NEB) overnight but not cleaned up. The homology arms were in 3:1 molar excess. 1.25 ug of total DNA was used (225 ng of upstream homology 626 bp, 225 ng of downstream homology 665 bp, and 800 ng of digested plasmid 4583 bp). Cells were streaked out from the −80C on YP+Glycerol. Four transformation cells were grown overnight in YPD, diluted into YPD, and allowed to grow for at least two doublings. We performed a Lithium Acetate transformation with 30 minutes at 30 C and 60 minutes at 42 C, followed by a two-hour recovery in synthetic dextrose minimal media without a nitrogen source, as recommended by Sasha Levy. We integrated plasmids in seven transformation batches, which were plated overnight on YPD and replica-plated onto YPD+G418 (200 ug/ml). Plates were stored at 4 C and then scraped with water, pooled, frozen into glycerol stocks, and mated. We collected an estimated 100,000 colonies, approximately fivefold coverage of the tiles. For 6/7 pools, we sequenced tiles before and after mating, finding that 67-97% of tiles were detected both before and after mating, indicating that the mating sometimes reduced library complexity.

### Yeast Mating

We mated each of the seven transformations independently to MY435 (FY5, MATalpha, YBR032w::P3 GFP ClonNat-R (pMVS102)). Downstream sequencing revealed that transformations with modest numbers of colonies (e.g. 4500) experienced no significant loss of complexity during mating, but transformations with more colonies (e.g. >20,000) experienced loss of complexity, up to 40% in one case. Subsequent matings were performed in larger volumes to avoid creating a bottleneck. Mated diploids were selected in liquid culture with YPD with 200 ug/ml G418 and 100 ug/ml ClonNat. After overnight selection, matings were concentrated and frozen as glycerol stocks.

### Cell Sorting

The day before sorting, a glycerol stock of mated cells (∼100 ul) was thawed into 5 mL SC+Glucose with 200 ug/ml G418 and 100 ug/ml ClonNat and grown overnight, shaking at 30 C. In the morning, the culture was diluted 1:5 into SC+Glucose with G418, ClonNat, and 10 uM ß-estradiol (Sigma). The culture was grown for 3.5-4 hours before sorting.

Cells were sorted on a BD Aria Fusion equipped with four Lasers (488 blue, 405 Violet, 561 Yellow-green and 640 Red) and eleven fluorescent detectors. We used two physical characteristics gates, first to enrich for live cells (FSC vs SSC) and second to enrich for single cells (FSC-Height vs FSC-Area). Cells were sorted by the GFP signal, the mCherry signal, or the ratio of GFP:mCherry signal. The ratio is a synthetic parameter that is very easy to saturate on the eighteen-bit scale available in the BD software. Great care was taken to change PMT voltage and the ratio scaling factor (5-10% depending on the day) to make the value of the top and bottom bins as different as possible. The dynamic range of our final estimate for activation domain activity is set by the value of the top and bottom bins. The maximum activation domain strength is 100% in the top bin, and assumes the value of the top bin. The minimum activation domain strength is 100% in the bottom bin and assumes the value of the bottom bin.

We performed our sorting experiment twice. In the first run, we pooled all of the transformants into one sample and sorted it by GFP/mCherry ratio, GFP-only, mCherry-only. We sorted one million cells per bin. For the ratio sort, we split the ratio histogram in eight approximately equal bins ^13^.

In the second round of sorting, we split the transformants into two pools, labeled A and B, so we could assess measurement reproducibility for independent transformants. Pool A and Pool B are true biological replicates. We sorted each pool by GFP/mCherry ratio, GFP-only, and mCherry-only. We used the comparison of the A and B pool measurements to assess measurement reproducibility of true biological replicates. We have never previously measured this biological reproducibility. On this day, we sorted 250000 cells per bin.

Sorted cells were grown overnight in SC-glucose. The next morning, gDNA was extracted with the Zymo YeaSTAR D2002 kit, using Protocol I with chloroform according to the manufacturer instructions. We have previously shown that growing cells overnight makes the gDNA extraction easier but does not change the computed activation domain activity ^13^.

### Amplicon Sequencing Library preparation

Amplicon sequencing libraries were prepared from genomic DNA in three steps. First, the general vicinity of the tile sequence was amplified with CP21.P14 and CP17.P12 using 100 ng of gDNA as template and yielding a 604 bp product that was cleaned up (Monarch PCR cleanup). In the second PCR, we added 1-4 bp of phasing on each end and the Illumina sequencing primer in 7-10 cycles with SL5.F[1-4] and SL5.R[1-3]. These seven phased primers were pooled and added to all samples. Four nanograms of the first PCR were used as template for the second PCR. Two microliters of the second PCR served as template for the third PCR. The third PCR added unique Index1 and Index2 sequences to each sample with an additional 7-10 cycles. These final products were cleaned up with PCR columns or magnetic beads (MacroLab at UC Berkeley) and submitted for sequencing. We performed 2×150 bp paired end sequencing in a shared Nova-Seq lane at the Washington University School of Medicine Genome Technology Access Center (GTAC). GTAC provided demultiplexed fastq files. We sequenced additional samples in shared Nova-seq lanes with MedGenome.

### Sequencing Analysis

After demultiplexing samples and pairing reads with PEAR, we kept only the reads where the tile DNA sequence contained a perfect match to a designed tile. For each eight bin sort, we performed two normalizations. We first normalized the reads by the total number of reads in each bin. Then, we normalized across the eight bins to calculate a relative abundance. We then converted relative abundances to an activity score for each tile by taking the dot product of the relative abundance with the median fluorescence value of each bin (**Table S8**). This weighted average is the measured activation domain activity. Tiles with fewer than forty-one reads were not included in the final dataset. These analysis scripts are available at github.com/staller-lab/labtools/tree/main/src/labtools/adtools. This preprocessing computed an activity for each tile in each experiment. Activity is uncorrelated with total reads (**Figure S2E**). The pooled ratio sort (BSY2) had 115.6 M reads. The Replicate A ratio sort had 934.5 M reads, and the Replicate B ratio sort had 697 M reads. Replicate A GFP had 33.1 M reads, Replicate B GFP had 31.6 M reads, Replicate A mCherry had 32.8 M reads, and Replicate B mCherry had 30.3 M reads.

### Measurement Reproducibility

We used the two measurements of independent transformants to assess the reproducibility of our measurements of true biological replicates (R = .870; **Figure S2A-D**). Reproducibility is higher (R= .919) for highly abundant tiles (>1000 reads).

We combined data from the two biological replicates. For tiles present in both populations (n= 11797), we averaged the two measurements and used the standard deviation as the error bar. For tiles present in only one population, we used that measurement and did not report error bars. These combined data agree very well with the pooled sort (R= .919; **Figure S2C**). Activity was saturated for forty-nine tiles, but most of these were measured with low fidelity because they had low read depth, and forty-seven were present in only one biological replicate. We identified forty-one tiles that were very highly active in both replicates and had high read depth in both replicates (**Table S11**). These we recommend for CRISPR Activation studies in yeast.

We assessed whether the mating introduced biological variability. We remated seven pools of the integrated library to the same reporter line, selected for diploids, pooled them, and resorted cells. This time we sorted 500,000 cells per bin. This measurement agreed with the initial experiments (R = 0.920; **Figure S2D**).

Inferred activity was not correlated with read count, which, as previously shown, is another indicator of high-quality data (**Figure S2E**).

We compared activity measurements to our previously published results ^13^. Previously, we used 44 AA regions, and here we used 40 AA tiles. We considered any 44 AA tile that contained one or our 40 AA tiles to be corresponding pairs. The extra 4 AA can modify activity, so the correspondence of these measurements will not be perfect. The observed Pearson correlation of 0.786 and Spearman correlation of 0.731 indicate the new data are of high quality and consistent with previous measurements (**Figure S2F**).

The technical reproducibility of our measurements at UC Berkeley are lower than the published reproducibility from sorting at Washington University in St. Louis ^13^. In both cases, we sorted the same cell population twice and created independent sequencing libraries. In 2018, the technical reproducibility was high, Pearson R = 0.988. The 2018 work had a smaller library (<5000 unique sequences) and sorted more cells (1-2 million cells per bin). Sorting more cells per library member increases the technical reproducibility of the measurement. The sorter operator in the 2018 work was more experienced than the sorter operator in this work (MVS), and the machine was maintained to a higher standard of operation, so the sorted populations were purer.

The eight bin ratio activity measurements are primarily driven by the GFP signal. Activity (ratio) is largely separable from abundance assessed by the mCherry sort (**Figure S3G-I**) and well-correlated with the GFP sort (**Figure S2J-L**).

### Determining a threshold for active tiles

The full distribution of tile activities has a peak at low activity, which, based on control sequences, is clearly inactive, with a heavy right shoulder and a heavy right tail. After trying many thresholds, we ultimately chose the top 20% (94,031) as a threshold for high activity.

To show that all the orthologs contain at least one active tile, we used multiple thresholds. At the stringent threshold, top 20%, there is only one ortholog with no active tiles, Canca1_23981 from *Tortispora caseinolytica*. This ortholog is an alternative (now deprecated) gene model for the Canca1_57326 protein, which contains an additional ninety-nine N-terminal residues with twenty-three overlapping active tiles that comprise two activation domains, the second of which overlaps the WxxLF motif. The short form of the protein starts at the WxxLF motif, further emphasizing that this motif is not sufficient to create an activation domain. Based on improved, transcript-based gene models, the short version, Canca1_23981, is likely a computational annotation error. There is more support for the long version, Canca1_57326. Given the weak evidence supporting the one potential exception, we conclude all of the Gcn4 orthologs are activators.

### Protein sequence parameters

We computed protein sequence parameters (Net charge, local net charge, Kyte Doolittle Hydrophobicity, Wimley White hydrophobicity, Kappa ^83^) with localCIDER ^40^. The OmegaWFYL_DE mixture parameter computes the mixture statistic between W,F,Y,L residues and D,E residues using the seq.get_kappa_X([‘D’,’E’],[‘W’,’F’,’Y’,’L’]) function in localCIDER ^41^. We predicted intrinsic disorder with MetaPredict2 ^84^. We counted motifs with regular expressions in Python with the “re” package.

The MAFFT algorithm aligns the WxxLF motif for all but three homologs. For three homologs, in the Full_length_homolog_dataframe, we corrected the “WxxLF motif location” parameter using the coordinates from the MSA. These species are the only ones outside the Ascomycota that have the motif. We suspect the WxxLF motif convergently evolved in these distance homologs because the context is very different and H rich.

Blastocladiomycota_jgi|Catan2|1097078|CE97078_6759, Blastocladiomycota_jgi|Catan2|1466814|fgenesh1_pg.199_#_9, and Blastoadioclmycota_jgi|Catan2|1506241|gm1.11555_g.

To predict helical propensity of homolog sequences, we used the Sparrow package in Python [https://github.com/idptools/sparrow]. A region was called helical if it contained five adjacent residues with over 50% chance of being helical. A large proportion of sequences have no residues with a >50% probability of being helical in this region. We consider this predictor to capture the propensity to form a helix in some context. To count proline residues in the region homologous to the known helix, we used the five AA upstream and five AA downstream of the WxxLF motif. From the 500 homologs in the MSA, there are 115 unique 15 AA regions around the WxxLF motif; twenty-three contain three prolines (20%) and three contain four prolines (2.6%).

Data were analyzed in Python with the matplotlib and seaborn packages.

### Imputing activity in the full-length homologs

We used the tile data to impute the activity of each position in each of the full-length homologs. The 19099 recovered tiles mapped to 68577 locations on the homologs (each tile matched to 3.6 homologs on average). We used a second-order Loess smoothing (twenty nearest points with the loess.loess_1d.loess_1d() function) across tiles to impute the activities of all positions in the 502 unique homologs. This quadratic smoothing can cause artifacts on the extreme ends of the protein, such as predicting negative activity. To remove this artifact, we constrained the imputed activity to be no more than the maximum measured and no less than the minimum measured in that homolog.

To validate the Loess smoothing, we averaged together all activities for all tiles that overlapped a position, equally weighing all tiles. These averages were more jagged because of the stepwise nature of the tiles. This simple average also created artifacts at the ends of the protein where only one tile is present. The Loess and average smoothing methods agreed well (97% had Pearson R > 0.80).

We used the imputed activities to create the heatmaps to visualize activity across the homologs. We tried many variations of these heatmaps but ultimately found that aligning the sequences on the start of the DBD or on the WxxLF motif was most informative. In the main text, we removed the six longest sequences to ease visualization.

We tested the hypothesis that insertions are enriched for active tiles by projecting activity onto the MSA. We defined insertions as the positions in the MSA with residues (non-gaps) in less than 1% of sequences (n < 5), which yielded 880/2690 (32.7%) of positions. In a two-sided t-test of the imputed activities of the insertion positions compared to all other positions, insertions were less active (p < 1e-135). We concluded that insertions are depleted for activation domain activity.

To estimate the activity at the WxxLF motif, we used the integral of the imputed activity from -10 to +10 around the W of the WxxLF motif. When this integral was below our activity threshold, we called sequences inactive in this region. Using this integral, 284 homologs (ninety-two unique sequences) had high activity (>150000) and twenty-seven homologs (thirteen unique sequences) had low activity, less than our activity threshold. Thirty-three had intermediate activity.

For motif enrichment, we performed a Welch’s t-test assuming unequal variances stats.ttest_ind(Sequences_WITH_Motif,Sequences_WITHOUT_Motif, equal_var=False).

To count activation domains on each TF, we combined active overlapping tiles, taking the union. With this method, we found 500 ADs with the WxxLF motif and 415 ADs without the WxxLF motif. As a result, we required more than forty residues between activation domains before they were called as two separate domains. Calling activation domains from the imputed activity map gives different results because some very close double peaks are split. With the smoothed data, there are 332 ADs with the WxxLF motif and 783 ADs without the WxxLF motif. The amino acid features of both classes are very similar.

### ANOVA

We used ordinary least squares regression (OLS) to create a baseline model for how composition controls activation domain function. We used ANOVA, OLS, and adjusted R-squared to compare models. See the Composition_ANOVA jupyter notebook for the full analysis. Briefly, we used the Python statsmodels ols(formula, ANOVA_DF).fit() function from the statsmodels package to fit the model, find coefficients, and compute adjusted R-squared values. We used the anova_lm(model, typ=2) function to find the sum of squares explained by each parameter. We used a Bonferroni multiple hypothesis correction to remove non-significant parameters and refit the model. In most cases, one iteration was sufficient to get a model where all parameters were significant. For the dipeptides, we used two interaction terms. All ANOVA parameters are in **Table S9**.

OLS regression on single amino acids explains 49.9% of variance in activity (**Table 1**, AUC = 0.9346, PRC = 0.7620, **Table S9**). Iteratively removing non-significant parameters led to sixteen residues, which explain 49.9% of variance. We repeated the regression with 400 dipeptides and found sixty-nine significant parameters that explain 60.2% of the variance in activity (**Table 1**, AUC = 0.9472, PRC = 0.8190). Half the variance in activity could be explained by composition alone, and dipeptides offered ∼10% improvement.

We predicted *de novo* motifs using the DREAM suite and then repeated the OLS ANOVA analysis using the motifs. We performed *de novo* motif searching on multiple slices of the data, but highly active (n=3524) vs. inactive (n=15575) were the most interpretable and gave the clearest signal in the ANOVA analysis. First, we ran the package STREME from the MEME suite to discover motifs that are enriched in a list of sequences relative to a user-provided control list.

For the OLS on *de novo* motifs, we used the motif counts provided by the DREAM motif prediction software (**Table S10**). For simplicity, in the parameter table, we refer to each motif as a string, but we used the PWM for actually finding motifs in each sequence with FIMO.

### Machine learning

We predicted activities on full length homologs using publicly available models, TADA, ADpred, and PADDLE^14,17,19,20^. All models were run on the SAVIO high performance computing cluster at UC Berkeley. TADA uses 40 AA windows, ADpred, 30 AA windows, and PADDLE 53 AA windows. For each TF, we tiled at 1 AA increments, spanning the full proteins (e.g. 1-40, 2-41 etc). For full-length TF analysis, we corrected the inferred activity at each position (Loess smoothing) with the predictions at each position. The smoothed data averages out some measurement noise so all the model performance is improved on smoothed data. For individual tile analysis, we used the center aligned score. We also tried maximum scores, average scores, and other variations, but chose center-aligned. ROC and PRC analyses were performed with the sklearn Python package.

Predicting the impact of mutating F residues in the central activation domains: we tile the 138 unique 70 AA central regions into 40 AA tiles spaced every 1 amino acid. For each tile, we computationally mutated each F individually, all pairs, all triplets, and all sets of four or more. For each mutant, we predicted activity. The mutants are predicted to have less activity. For each mutant, we also computed the change in activity. Finally, we grouped the changes in activity based on the conservation of each F residue.

### Datafiles

All the raw sequencing data has been deposited at NIH SRA Accession #PRJNA1186961: http://www.ncbi.nlm.nih.gov/bioproject/1186961

All the analysis scripts are deposited on github and Zenodo:

10.5281/zenodo.14201918

https://github.com/staller-lab/Gcn4-evolution

github.com/staller-lab/labtools/tree/main/src/labtools/adtools

https://github.com/staller-lab/Gcn4-evolution

All the processed data are attached in supplemental tables (**Tables S5 - S7**).

Processed sequencing read counts are in **Table S13**.

The ‘masterDF’ dataframe contains each designed tile (**Table S5**). Tiles that were not measured have activity recorded as nan or 0. The ‘orthorlogDF’ dataframe contains all tiles associated with each original full-length homolog (**Table S6**). As a result, tiles occur multiple times because they map to multiple homologs. The ‘NativeLocation’ is the position of the tile relative to the first amino acid. The ‘NormLocation’ is the position of the tile relative to the WxxLF motif. Finally, the ‘FullOrthoDF’ dataframe contains one entry for each full-length homolog, and each column contains an array with values for each position (**Table S7**), such as imputed activity at each position and local charge from localCIDER. The location of the bZIP DNA-binding domain was identified with the <u>InterPro signature (IPR004827)</u>.

## Acknowledgments

We would like to thank Nick Ingolia, Zeba Wunderlich, Rachel Brem, Alex Holehouse, Andrew Murray, Shahar Sukenik, Michael Botchen, and Ashley Wolf for helpful comments on the manuscript. Sumanth Mutte for finding the initial homologs. We thank Lucia Strader, Nicholas Morffy, Ross Sozzani, Lisa Van den Broeck, Hunter Nisonoff, and Jennifer Listgarten for helpful discussions. Nick Morffy and Lucia Strader for the yeast genome targeting plasmids. Igor Grigoriev identified the deprecated *Tortispora caseinolytica* gene models. Weijing Tang performed exploratory analyses not included in the final manuscript. The Regents of the University of California have filed an invention disclosure based on the findings of this study. The DHY213 BY superhost strain used for library construction was a generous gift from Angela Chu and Joe Horecka, and requests for this strain should be directed to them.

