## Supplemental Figures for "Evolutionary turnover of key amino acids explains conservation of function without conservation of sequence in transcriptional activation domains"

Figure S1: Overview of Gcn4 homolog selection
Figure S2: Measurement quality and reproducibility
Figure S3: Control activation domains and tiles from DBDs
Figure S4: Heatmap of homolog activity projected onto the species tree
Figure S5: Projecting the heatmap of activity onto the MSA
Figure S6: Zooming into active regions in the MSA
Figure S7: The 138 unique central activation domain sequences are diverse.
Figure S8: The Gcn4 ortholog dataset efficiently identified key sequence features controlling activity
Figure S9: Slicing through the 2-D landscape
Figure S10: Comparison of amino acid turnover in most active homologs compared to all homologs
Figure S11: Conservation of key residues in other activation domains mirror Gcn4
Figure S12: Neural network models for predicting activation domains from amino acid sequence perform well on the Gcn4 homologs
Figure S13: A key coactivator of Gcn4, Med15/Gal11 shows high conservation
Figure S14: Predicting tile binding to Med15/Gal11 with FINCHES
List of Supplemental Tables
References

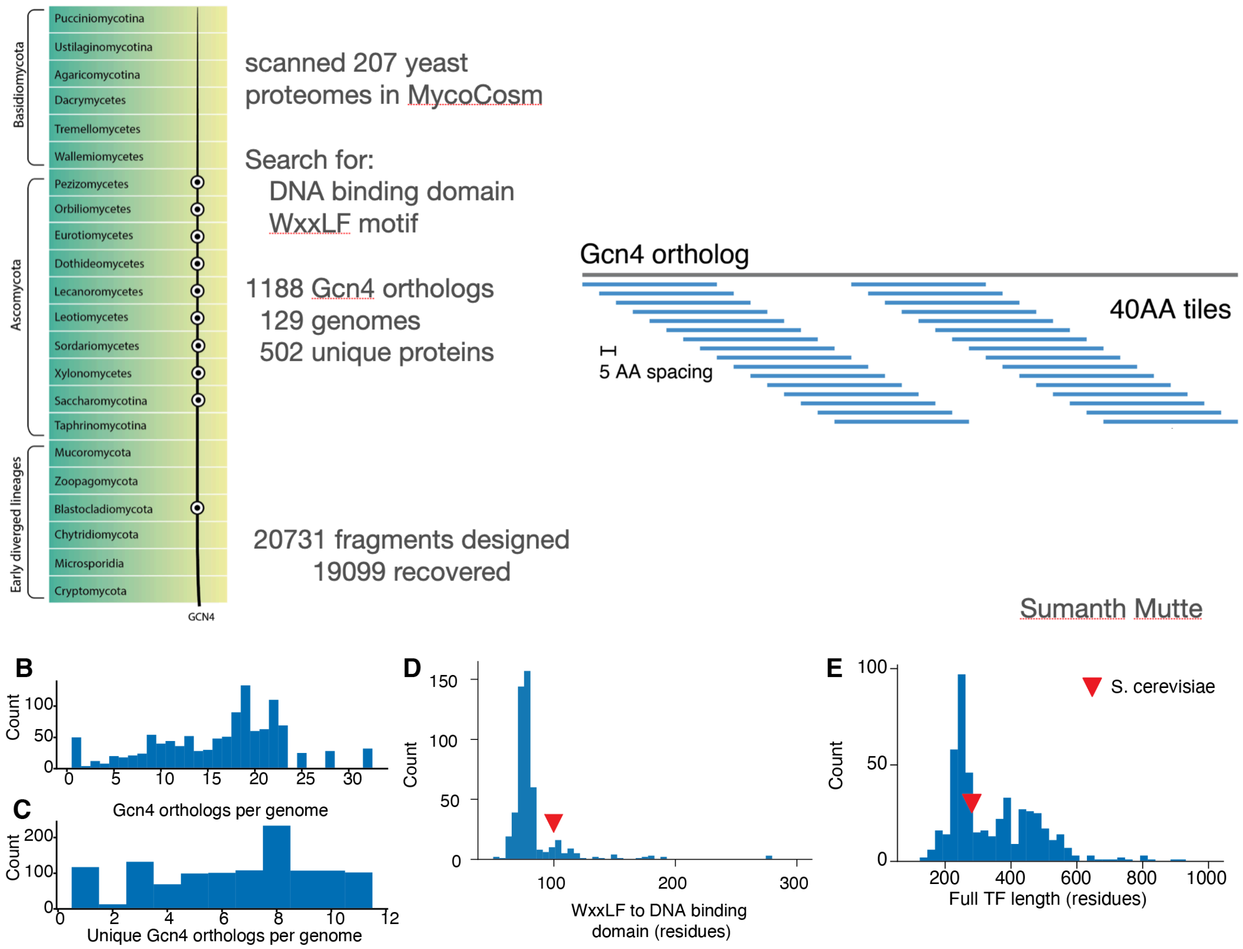


**Figure S1: Overview of Gcn4 homolog selection**

A) 207 diverse fungal proteomes were selected to represent the diversity of the Mycosm database. The selected proteomes came from the subdivisions indicated with circles. We scanned the proteomes for Gcn4 homologs using the DNA binding domain and the WxxLF motif. For the DNA binding domain, we used the IPR004827 profile HMM from Interpro. The WxxLF motif has been shown to be conserved [1,2]s and we used the regular expression Wx[SPA]LF. Each of the 502 unique homologs we identified was tiled into 40aa overlapping fragments. 19099 designed fragments were detected after yeast transformation and 18947 passed abundance thresholds to be included in the analysis.

B) Histogram of homologs identified in each genome.

C) Histogram of unique homologs identified in each genome. Multiple gene models or splice forms can yield the same protein sequence.

D) Distribution of distances from WxxLF motif to the start of the protein.

E) The distribution of homolog lengths varies considerably.

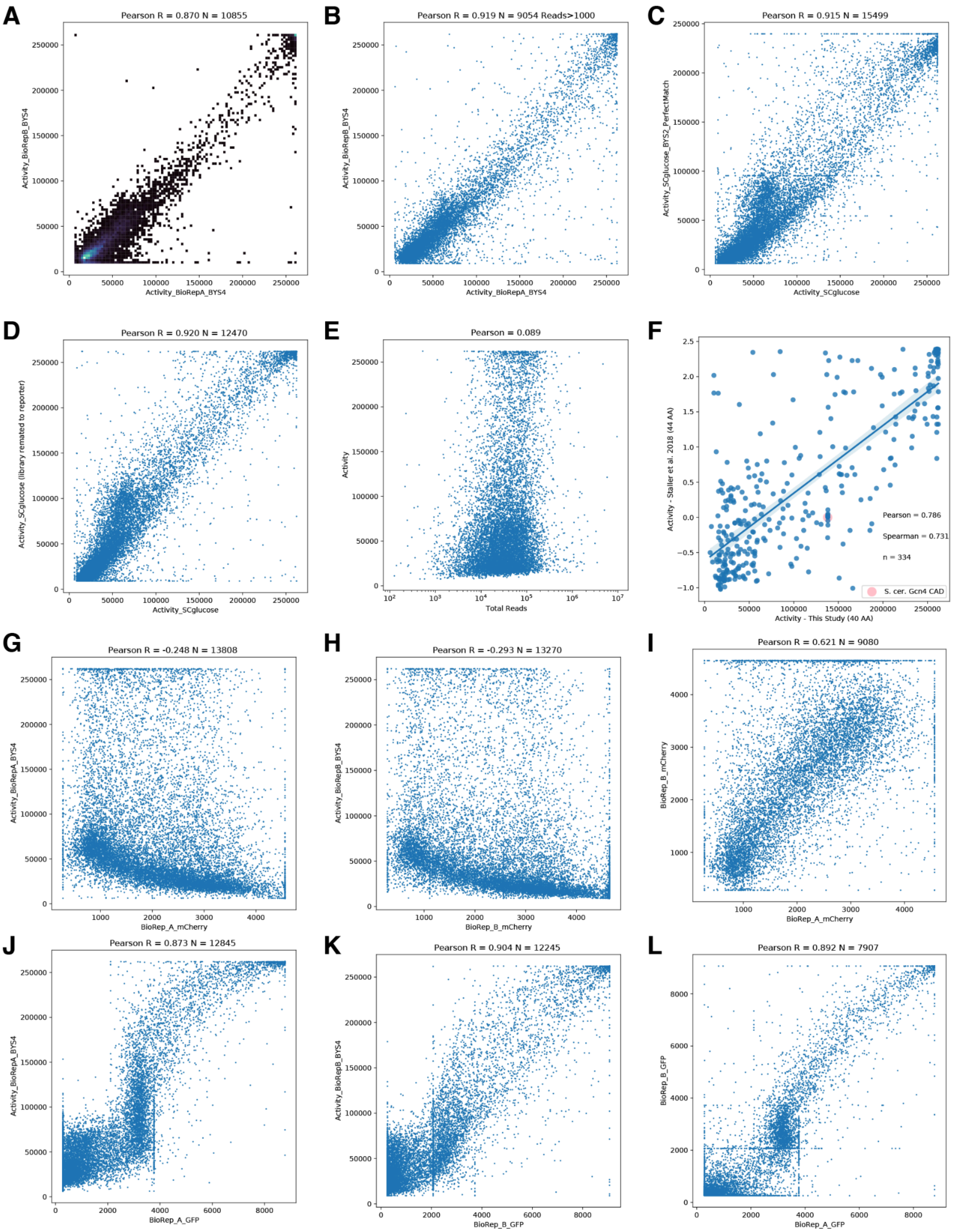


**Figure S2: Measurement quality and reproducibility**

A) Measurement reproducibility when we integrated the plasmid library into yeast in Pool A and Pool B. These are independent biological replicates. This panel shows all the data. Color indicates the number points (tiles) in each pixel.
B) Filtering out tiles with fewer than 1000 reads improved measurement reproducibility.
C) For the main dataset, we combined the two biological replicates. This combined number was well correlated with a separate experiment in which we physically mixed all the yeast together after integration, induced, and sorted cells.
D) Measurements of two independent matings of the same library of integrated synthetic TFs. This captures biological variation from mating and technical variation from two independent sorts months apart.
E) Activity is uncorrelated with total read count, a proxy for abundance in yeast.
F) 40AA tiles in this work were compared with 44AA tiles in a previous study[3]. The measurements generally agree.
G) The Activity (GFP/mcherry ratio) is largely separable from abundance (mCherry). Pool A data.
H) Similar to G with Pool B.
I) mCherry measurement reproducibility. Vertical lines arise from tiles that have low abundance in one replicate and are only found in one sorting bin.
J) The GFP signal is consistent with the Activity (GFP/mCherry ratio) in Pool A.
K) Similar to J with Pool B.
L) GFP measurement reproducibility. There are four bins.

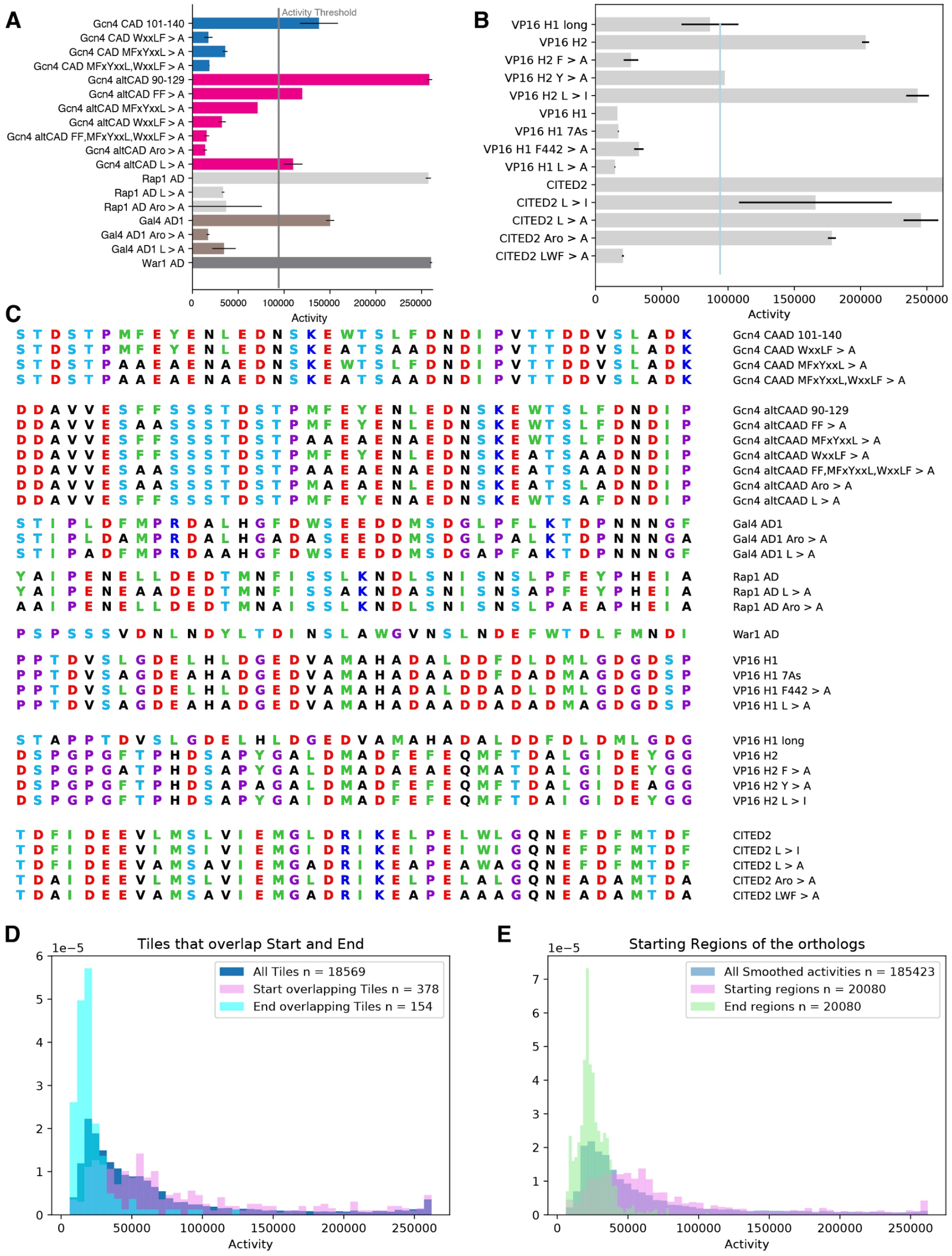


### Figure S3: Control activation domains and tiles from DBDs

A) Activity of control activation domains from yeast TFs and hand designed mutants.

B) Control activation domains from human and human viruses. Aromatic residues make larger contributions to activity than leucine residues.

C) Sequences of the hand designed mutations.

D) Tiles from the C-terminus (End tiles) that overlap the DBD have low activity in the assay. Tiles from the N-terminus (Start tiles) have activity that matches the full distribution.

E) A similar analysis using the imputed, smoothed activities at each position. The first 40 and last 40 residues are used for each analysis. End regions that overlap the DBD have little-to-no activity. Start regions from the N-terminus resemble the full distribution.

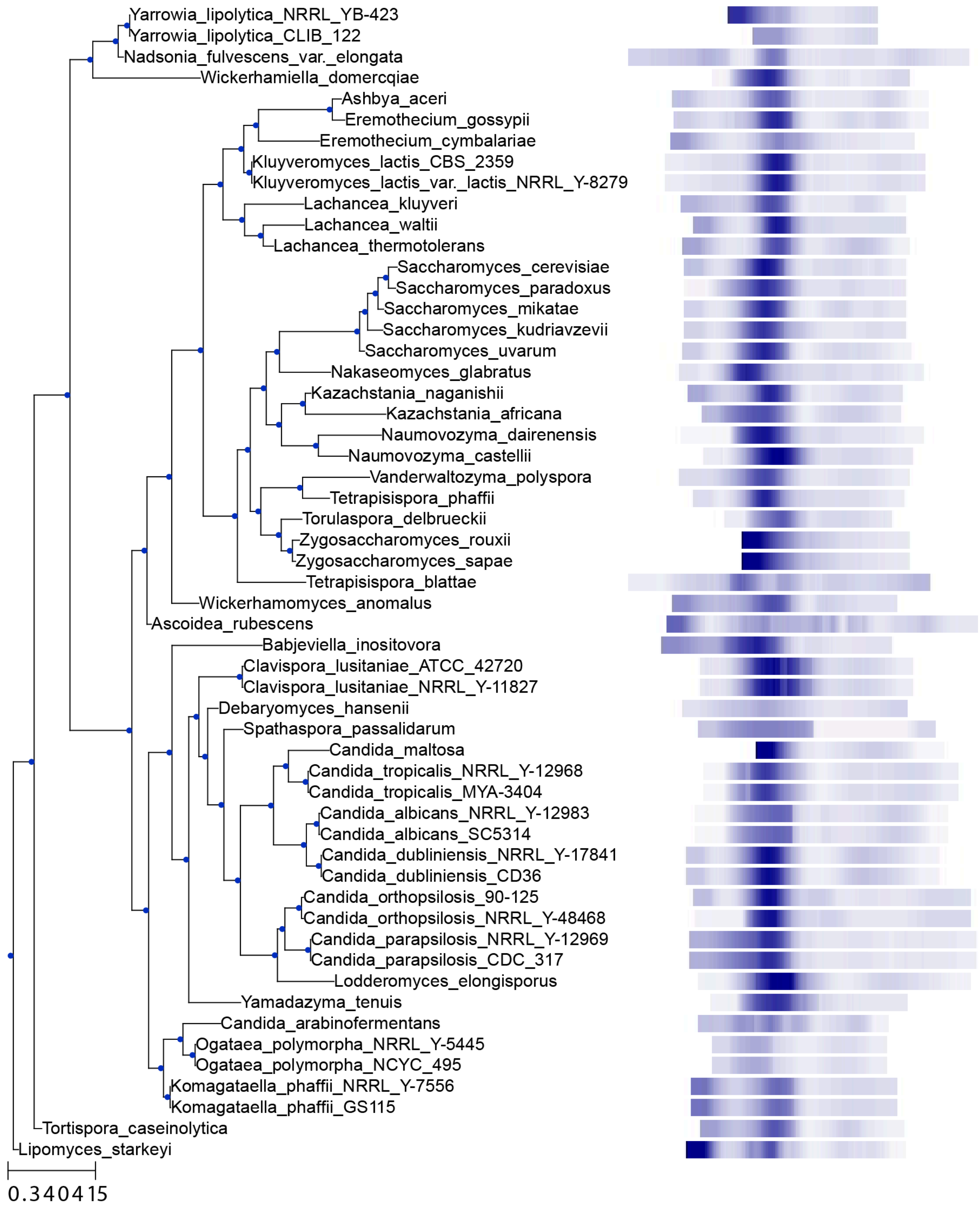


**Figure S4: Heatmap of homolog activity projected onto the species tree**

For our homologs that were part of the Y1000+ project, we visualized heatmaps of the activity on the species tree. Sequences are aligned on the WxxLF motif. Scale bar from [4]. Most of the species close to *S. cerevisiae* have one activation domain, but it can move around. Tree visualization was made with ETEToolkit [5].

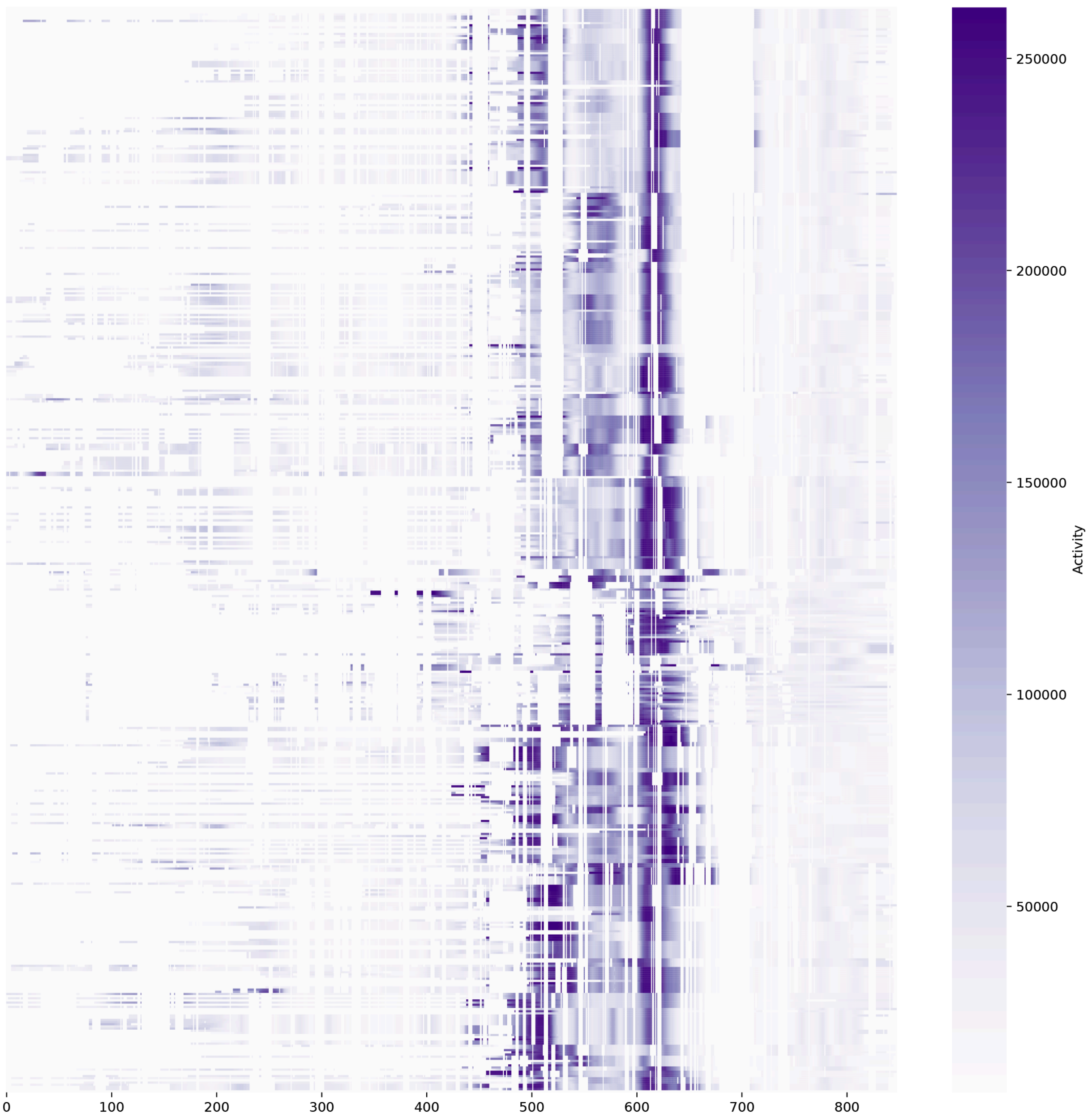


**Figure S5: Projecting the heatmap of activity onto the MSA**
Inferred activity at each position was projected onto the MSA of 500 homologs. Columns in the MSA with fewer than 25 sequences (5%) were filtered out. The WxxLF motif is at approximately position 625.

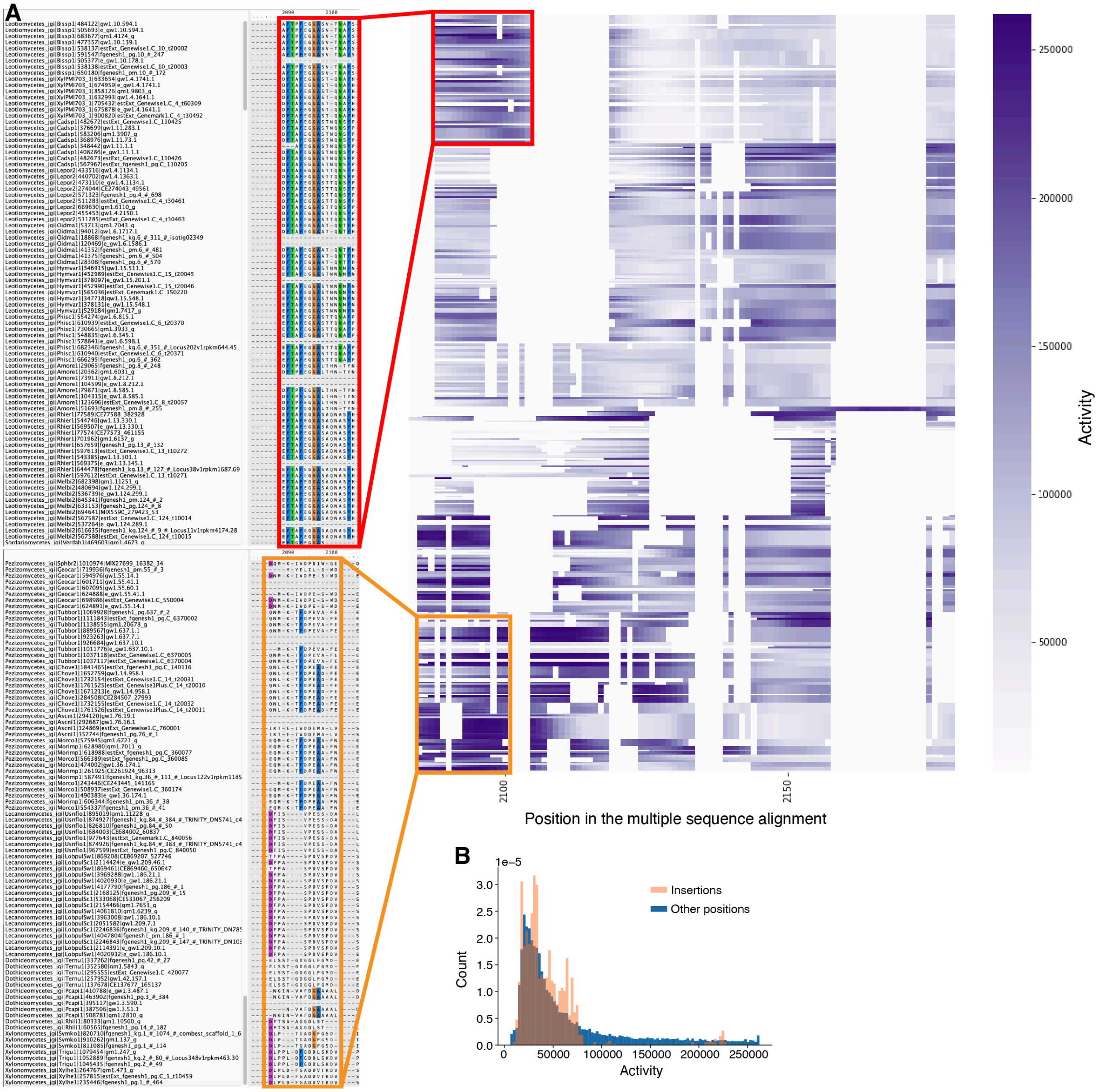


**Figure S6: Zooming into active regions in the MSA**

A) This zoomed inset of the MSA shows how N-terminal activation domains are mildly conserved. The boxed N-terminal activation domain appears to be conserved between two distant clades. Close examination reveals the sequences are highly diverged. If these are descendants from the same ancestral activation domain, they have diverged beyond recognition. Majority consensus positions are highlighted in color. Note, all columns are included in this version. This region also contains a rare insertion with high activity (right).

B) The histogram shows that insertions are depleted for activity (p = 2.397 e-136, uncorrected one sided t-test).

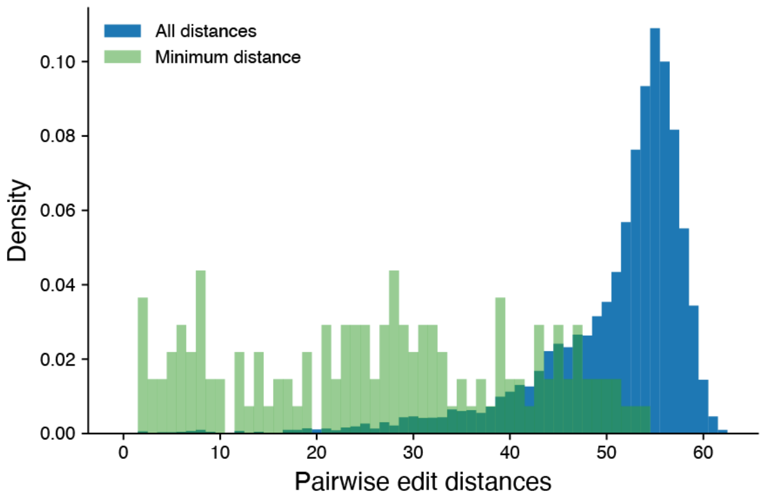


**Figure S7: The 138 unique central activation domain sequences are diverse.**
For all pairs of the 138 unique central activation domain regions (-50 and +20 from the WxxLF motif), we computed the pairwise edit distance (blue). For each sequence, we recorded the minimum distance to the most similar sequence (green). The maximum possible distance is 67 because the sequences are 70 AA long and they are all aligned on the WxxLF motif.

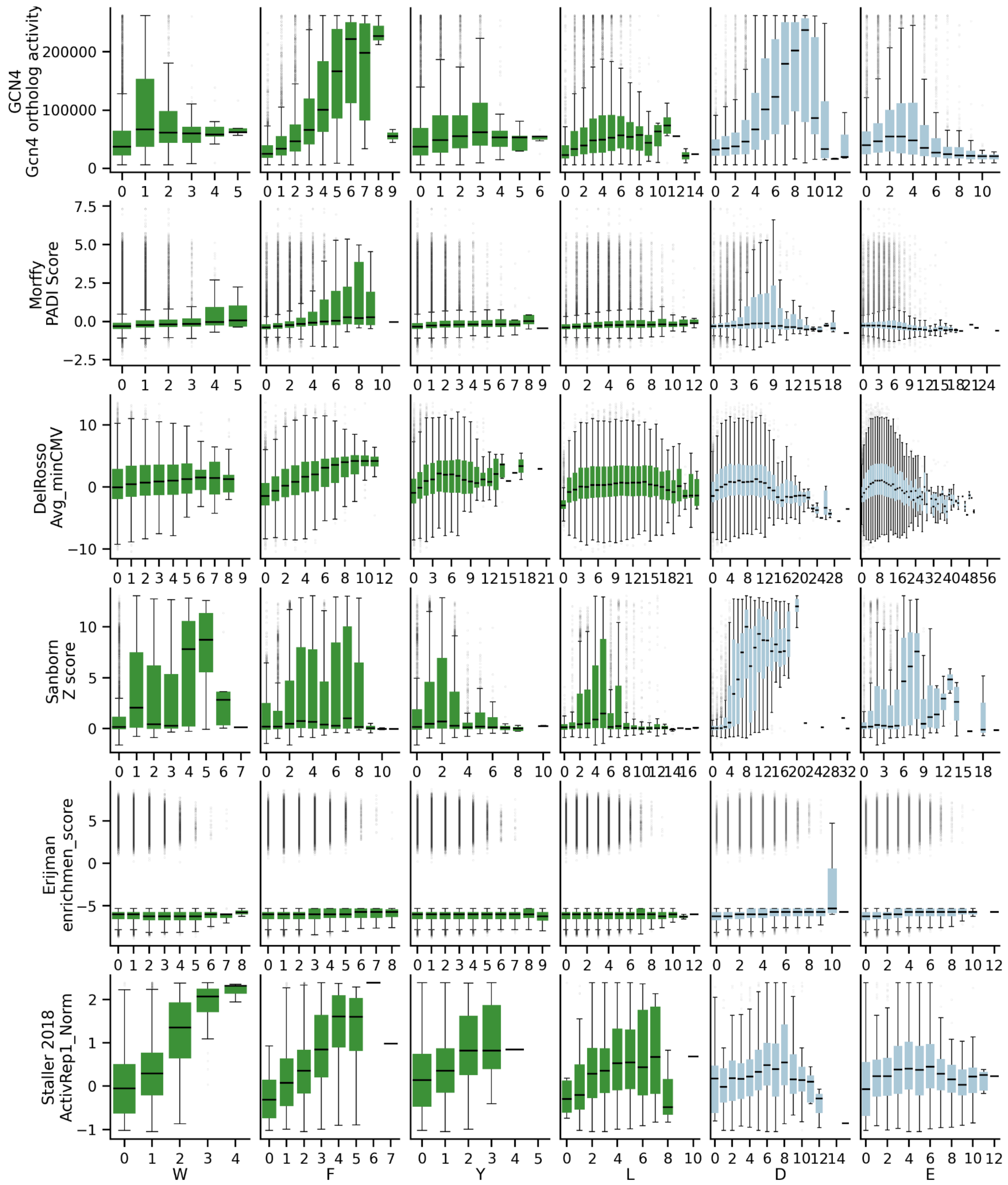


**Figure S8: The Gcn4 ortholog dataset efficiently identified key sequence features controlling activity**

We compared the Gcn4 ortholog dataset to other published high-throughput datasets. Many of the signals for sequence features that control activity are more visible in the Gcn4 ortholog dataset than in previous datasets. For example, the difference between D and E or the effect of F. Souce data: GCN4: this study (length 40). Morffy: All sequences (length 40). DelRosso: CRTF tiling library and AD mutants library (most length 80, some 70). PADDLE: TF tiles, AD mutants, and 53 aa mutants (length 53). Erijman: All sequences (length 30). Staller 2018: “ActivityCompleteMedia Replicate1_Normalized” (length 44).

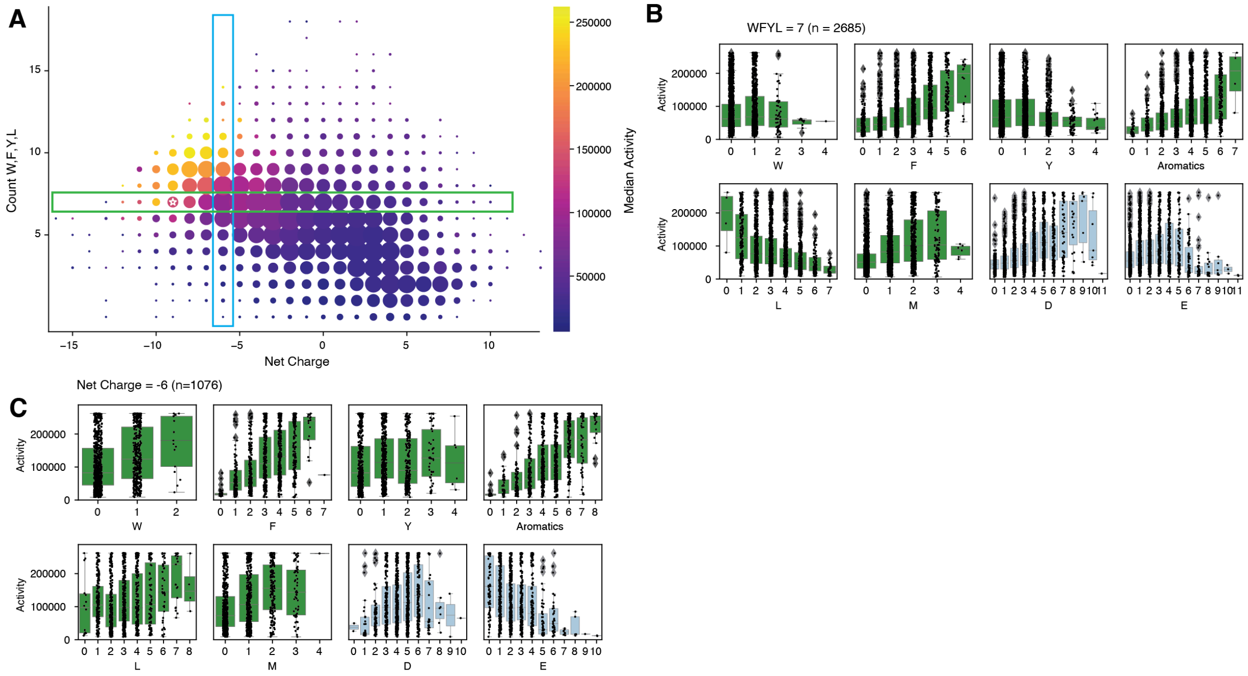


**Figure S9: Slicing through the 2-D landscape**

A) Similar to Figure 4A. Tiles with WFYL=7 are shown in the green box. Tiles with net charge=-6 are shown in blue box.

B) Box plots for the tiles with WFYL = 7, green box in A. In this set, the difference between D and E is very visible.

C) Box plots for all the tiles with net charge = -6. In this set, it is clear that L and M play a supporting role, boosting activity.

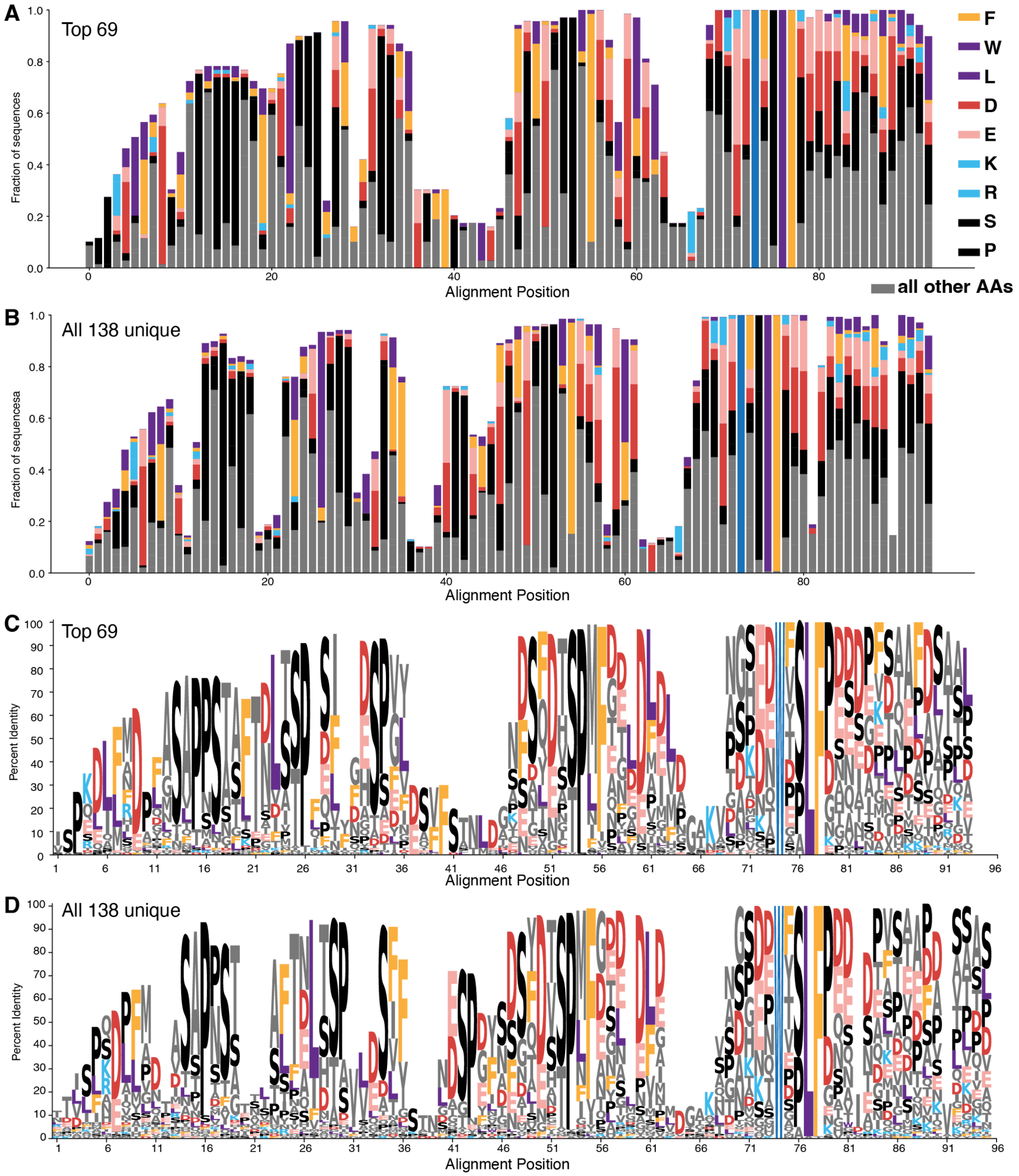


**Figure S10: Comparison of amino acid turnover in most active homologs compared to all homologs**

A) Stacked barplot of amino acid conservation in the 69 most active homologs. Similar toFigure 5A, but showing all residues.

B) Stacked barplot for all 138 unique homologs.

C) Sequence logo of MSA of sequences in A. Reproduction of Figure 5B.

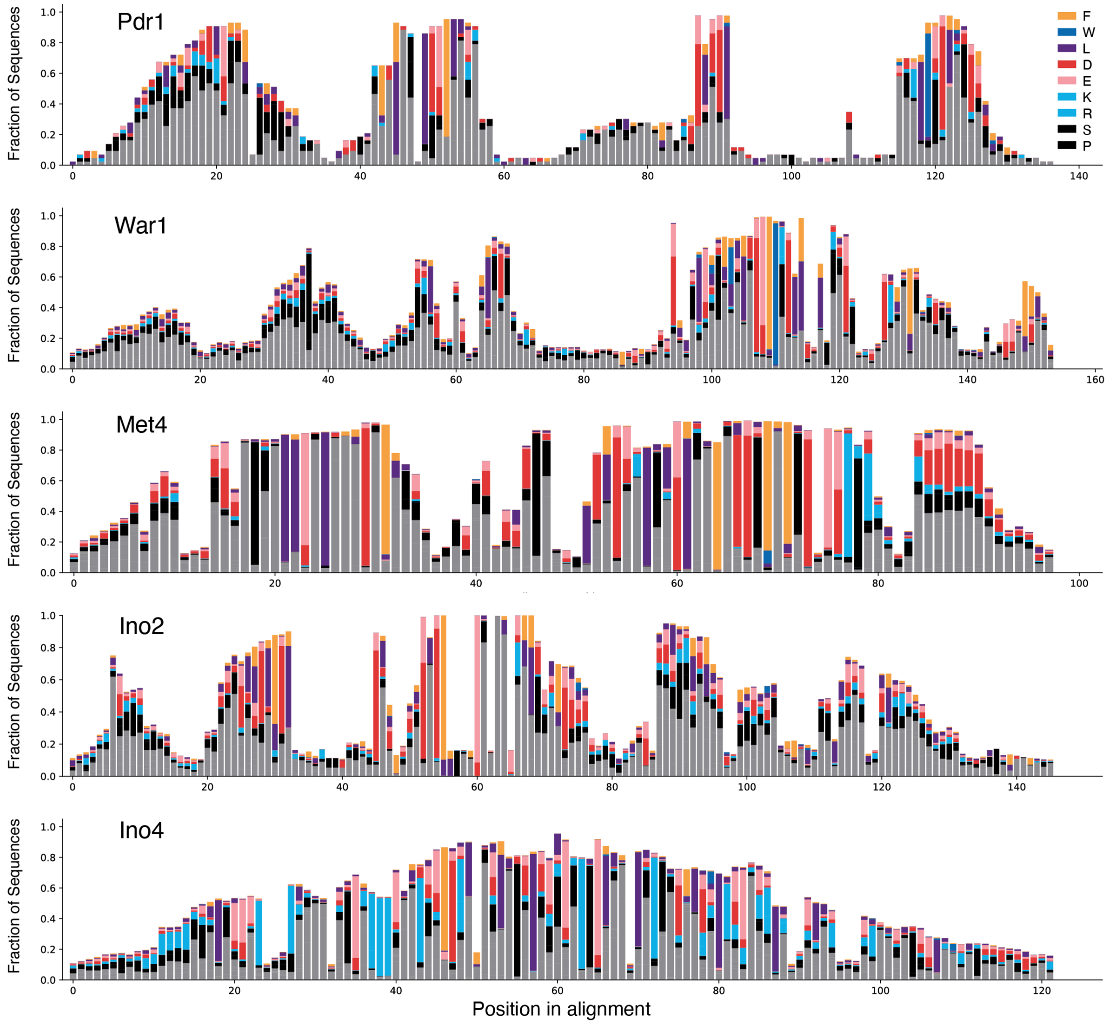


**Figure S11: Conservation of key residues in other activation domains mirror Gcn4**
For Pdr1, we took active sequences from Sanborn et al. 2021. For the other activation domains, we searched for homologs in the Y100+ collection using HMMER. The profile hidden Markov model was generated using the Yeast Genome Order Browser alignment. homologous sequences were further filtered to ensure they contained the DNA binding domain. For all homologous sequences, we also performed a blast search against the *Saccharomyces cerevisiae* genome and filtered out those that did not have the TF of interest as the top hit. For War1, two additional *S. cerevisiae* TFs appeared in the HMMER search results. Therefore, we performed additional HMMER searches for both TFs and filtered out sequences that had higher scores in the other TF HMMER searches compared to those of the War1 search. All the major patterns for Gcn4 are apparent in these regions. The acidic residues interconvert. Aromatic residues are highly conserved (Met4) and can turnover (War1). There is some interconversion between F and L in Ino2. In War1, in the FWxxLF motif, only the FW is conserved. Prd1 n = 43, War1 n=1082, Met4 n=922 , Ino2 n = 221, Ino4 n= 1419.

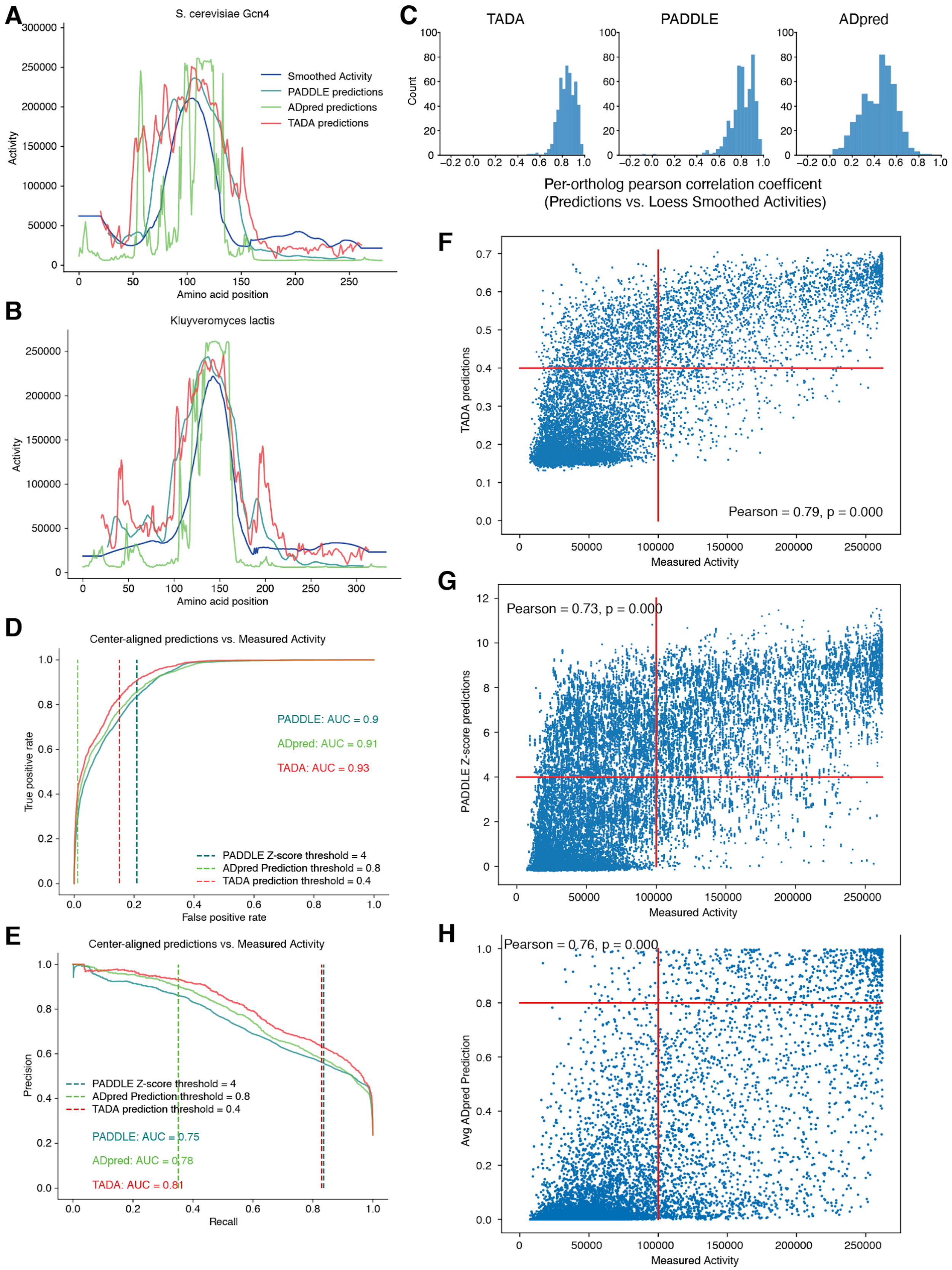
A
S. cerevisiae Gcn4
Smoothed Activity
PADDLE predictions
ADpred predictions
TADA predictions
Activity
Amino acid position
B
Kluyveromyces lactis
Activity
Amino acid position
C
TADA
PADDLE
ADpred
Count
Per-ortholog pearson correlation coefficent
(Predictions vs. Loess Smoothed Activities)
D
Center-aligned predictions vs. Measured Activity
PADDLE: AUC = 0.9
ADpred: AUC = 0.91
TADA: AUC = 0.93
PADDLE Z-score threshold = 4
ADpred Prediction threshold = 0.8
TADA prediction threshold = 0.4
True positive rate
False positive rate
E
Center-aligned predictions vs. Measured Activity
PADDLE Z-score threshold = 4
ADpred Prediction threshold = 0.8
TADA prediction threshold = 0.4
PADDLE: AUC = 0.75
ADpred: AUC = 0.78
TADA: AUC = 0.81
Precision
Recall
F
TADA predictions
Pearson = 0.79, p = 0.000
Measured Activity
G
Pearson = 0.73, p = 0.000
PADDLE Z-score predictions
Measured Activity
H
Pearson = 0.76, p = 0.000
Avg ADpred Prediction
Measured Activity

**Figure S12: Neural network models for predicting activation domains from amino acid sequence perform well on the Gcn4 homologs**

A-B) Measured activity and predicted activity (3 models) for *S. cerevisiae* and *K. lactis*. The models approximate the location of the activation domain reasonably well.

C) For each full length TF, we correlated smoothed measured activity with the predictions. The models can approximate the general location of activation domains. This comparison underweights errors in estimating activation domain boundaries. TADA performs better than the first generation models.

D) Receiver Operator Characteristic (ROC) curves for model performance on individual tiles. For each tile, we used the center aligned predicted activity because the predictors use windows of different lengths. TADA outperformed the older models.

E) Precision Recall curve (PRC) for model performance on individual tiles. TADA outperformed the older models.

F-H) Scatter plot for measured activity and predicted activity of each tile using center aligned data. Vertical red line, 100,000 activity units to guide the eye, slightly higher than the top 20% active threshold. Horizontal red lines, the predictor activity thresholds recommended by the authors of each model. Performance on individual tiles is worse than performance on full length TFs.

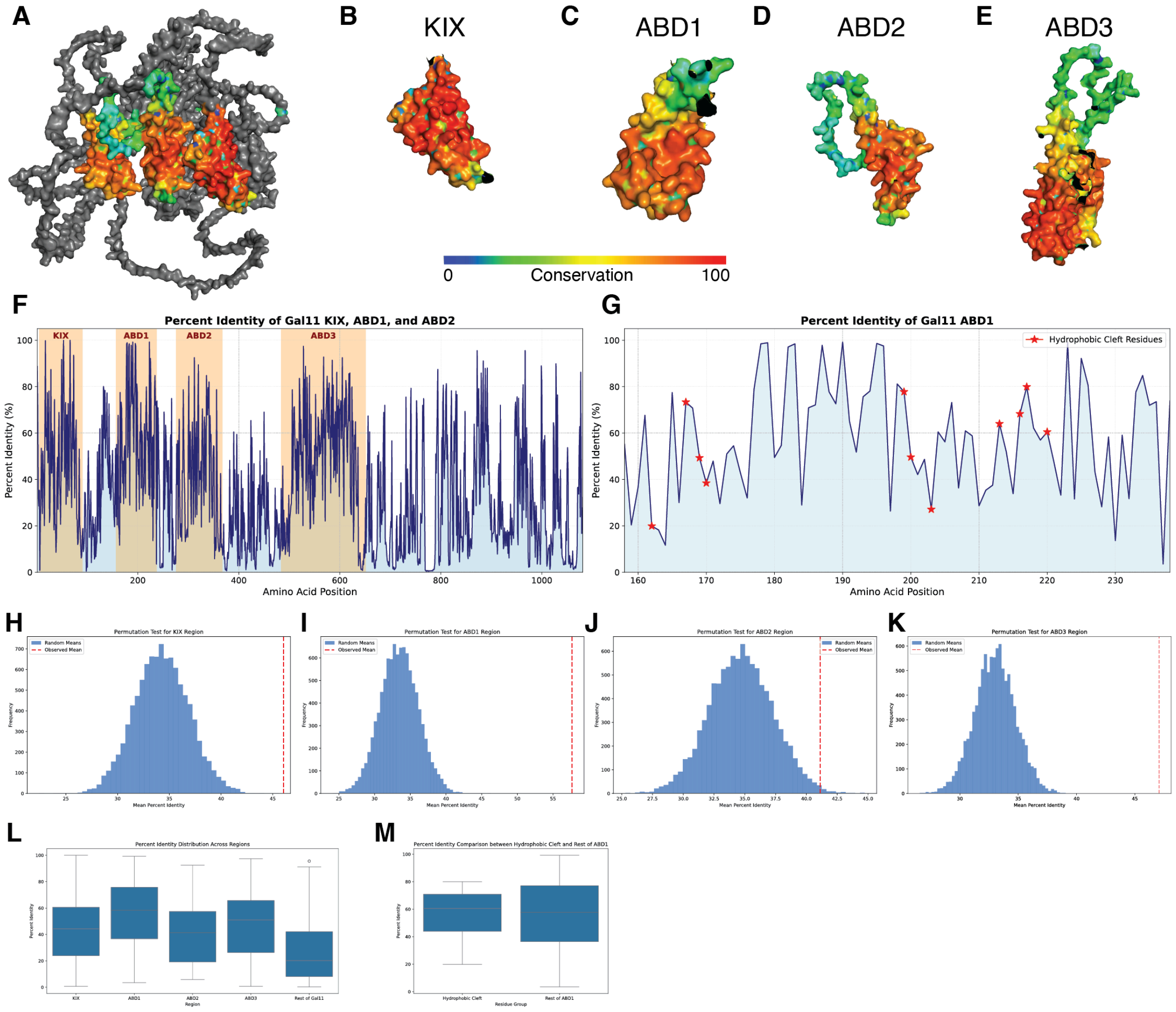


**Figure S13: A key coactivator of Gcn4, Med15/Gal11 shows high conservation**

A) We collected 653 Med15 sequences from the Y1000+ collection and created an MSA. The AlphaFold structure of *S. cerevisiae* Med15/Gal11 with the KIX, ABD1 and ABD2 domains colored by their conservation (percent identity) in the MSA.

B-E) The KIX, ABD1, ABD2, and ABD3 domains colored by conservation in the MSA.

F) The conservation profile of the full protein with domains highlighted (Gaps trimmed).

G)The hydrophobic residues of ABD1 that engage with Gcn4 [6].

H-K) Permutation tests for percent identity of each domain compared to the rest of the protein.

L) Distributions of conservation in each domain.

M) The residues that make contact with Gcn4 in ABD1 are not more significantly conserved than those of the rest of the domain. Using only the YGOB high quality homologs gave similar results for all panels except for M, where there was more conservation of the contacting residues.

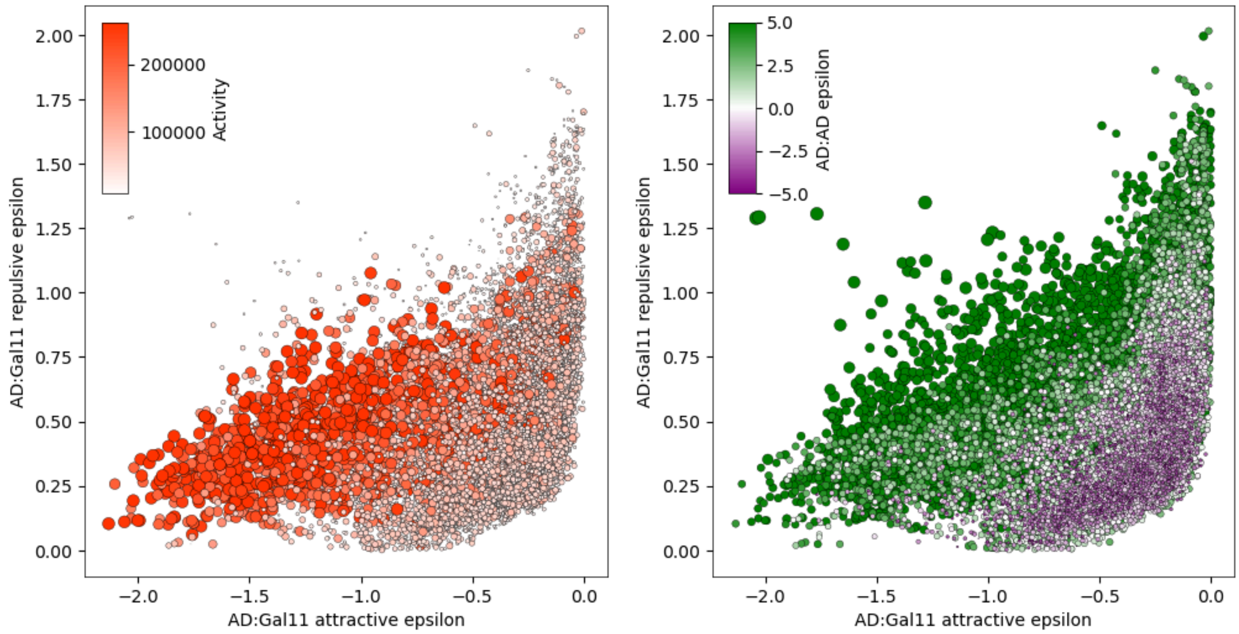


**Figure S14: Predicting tile binding to Med15/Gal11 with FINCHES**

FINCHES is a computational method for predicting binding between an IDR and a folded domain. During its initial development, it was benchmarked against activation domain binding to Med15/Gal11[7,8]. Running FINCHES on the homolog tiles predicts that binding to Med15/Gal11 is the primary molecular mechanism for activation by the Gcn4 homologs. The most active tiles have high attraction to Med15/Gal11 and low repulsion from Med15/Gal11[7].

### List of Supplemental Tables

1. Table_S1_Unique Full length homologs
2. Table_S2_MAFT multiple sequence alignment of 500 homologs
3. Table_S3_Control_Sequences
4. Table_S4_Primer_Sequences
5. Table_S5_Tile_Activities_Properties_Dataframe (masterDF)
6. Table_S6_homolog_Tile_dataframe (homolog DF)
7. Table_S7_Full_length_homolog_dataframe (FullLenthOrthoDF)
8. Table_S8_Gcn4 homologs FACS summary
9. Table_S9_ANOVA parameters
10. Table_S10_De novo found motifs
11. Table_S11_VeryStrongADsWithHighReproducibility
12. Table_S12_GCN4_homologs_bzip_DBD_Locations
13. Table_S13_Processed_Read_Counts
14. Table_S14_YGOB_SelectionCoefficients
